# A Requirement for Partial Dehydration of the K^+^ Ion Governs Gating of the Shaker K^+^ Channel: Quantum Calculations Show Complex Interactions of Ions, Water, Protons, and Protein Side Chains

**DOI:** 10.64898/2026.07.01.735716

**Authors:** Alisher M. Kariev, Regina R. Monaco, Michael E Green

## Abstract

There is a vast literature on the voltage gating of ion channels, with a fairly large fraction concerned with potassium channels, especially of the K_V_1 family, including *Shaker*. Experimental evidence derived from protein structure has been interpreted to give gating mechanisms that largely disregard water. We propose that the K^+^ ion, in order to pass through the gating region and enter the cavity pore, must be largely dehydrated. Competitive interactions of single hydration shell water molecules at the gate, with counterions, protein, or other water molecules, can remove one water at a time; not every water molecule is removed. There are several such interactions for the ion hydration shell; for the ion to pass through the gating region, there must be enough such interactions to leave the ion with at most two hydrating water molecules, in which case the gate is open. Protein conformational changes are small, and mostly unimportant. The hypothesis has a second part: protons, previously shown to be candidate carriers of the gating current (Kariev and Green, JPC B, 2019, Membranes, 2022, 2024) are capable of reaching the gate; adding four protons to the gate prevents dehydration, leaving the ion with at least three hydrating water molecules, enough to block passage. Quantum calculations presented here support the dehydration part of the hypothesis; they also mostly support the second part, concerning the protons, but further work will be required to fully confirm this. The hypothesis explains the experimental finding that the P475D mutant is essentially constitutively open, while the P475S mutant, with a wider gate opening, is closed at all relevant potentials; the computations presented here show the mechanism for this in detail, further confirming the first part of the hypothesis, and largely but not completely confirming the second part, concerning protons, while showing where further work is needed. This mechanism can also qualitatively account for flicker noise and fluctuations, and their consequences.

## Introduction

There is a vast literature on ion channels, the proteins that allow ions, particularly K^+^, Na^+^, and Ca^2+^, in and out of cells. Of this, a good fraction concerns voltage gated ion channels (VGIC) such as K_V_1.2 or *Shaker*; many consider gating methods or mechanisms (1–5) (6–9); this brief list of references is far from comprehensive. There exists a structural biology basis for discussion of the gating mechanisms, but with disagreement on significant details. However, the structures are almost all of the open state, although now there are structures of closed states beginning to appear (6, 10), even if these have been determined for potassium channels other than K_V_1.2 or *Shaker*. The channels in question have a structure consisting of four voltage sensing domains (VSD) with four transmembrane segments each, each domain apparently attached through a linker to a pore segment through which the ion passes. The reason the closed channel has proved far more challenging is that getting a structure with voltage applied is difficult. The general view of a closed structure is that one of the transmembrane segments from each VSD moves, some connection is made to the linker, an intramolecular segment between VSD and gate, which moves to exert pressure on the protein at the pore opening, and as a consequence there is mechanical closure of the space at the gate entrance. We have earlier reviewed this literature in detail (11), showing reasons to question this mechanism.

> Definitions of “top”, “bottom” and related words for figures and text: All figures show the axis that defines the approximate path of the permeant K^+^ oriented vertically. The intracellular end of the axis is shown at the bottom of all figures, while the upper end of the figures is closest to the entrance of the to the pore cavity. This convention is used throughout the paper, including in the text. Wherever the words “upper” and “top” are used, they indicate the region nearest the entrance to the pore cavity, while “lower” and “bottom” are used to indicate the part of the gating region nearest the intracellular opening. Thus the words “upper”, “top”, “lower”, “bottom”, are defined to indicate direction or location in the gate region.

While it is necessary to have the protein structure of the channel, this cannot be the full story. The gate region, as shown by most of these references, has an intracellular opening of around 15-17 Å. The opening tapers to roughly 12 Å about 6 Å further from the intracellular opening, which generally has a serine, and, closer to the center of the pore, an asparagine or glutamine (Fig. 1). As the gate is not perfectly symmetric, these dimensions are not precisely defined. At the upper end of the region there is a ring of prolines (connected to each other through valines) and the entrance to a cavity that connects to the selectivity filter at the outer end of the pore; this pore cavity contains water that can (partially) rehydrate the ion: see Figure 1 for the architecture of the gate protein that provides the framework for the hydration and dehydration of the ions.

**Fig. 1:**
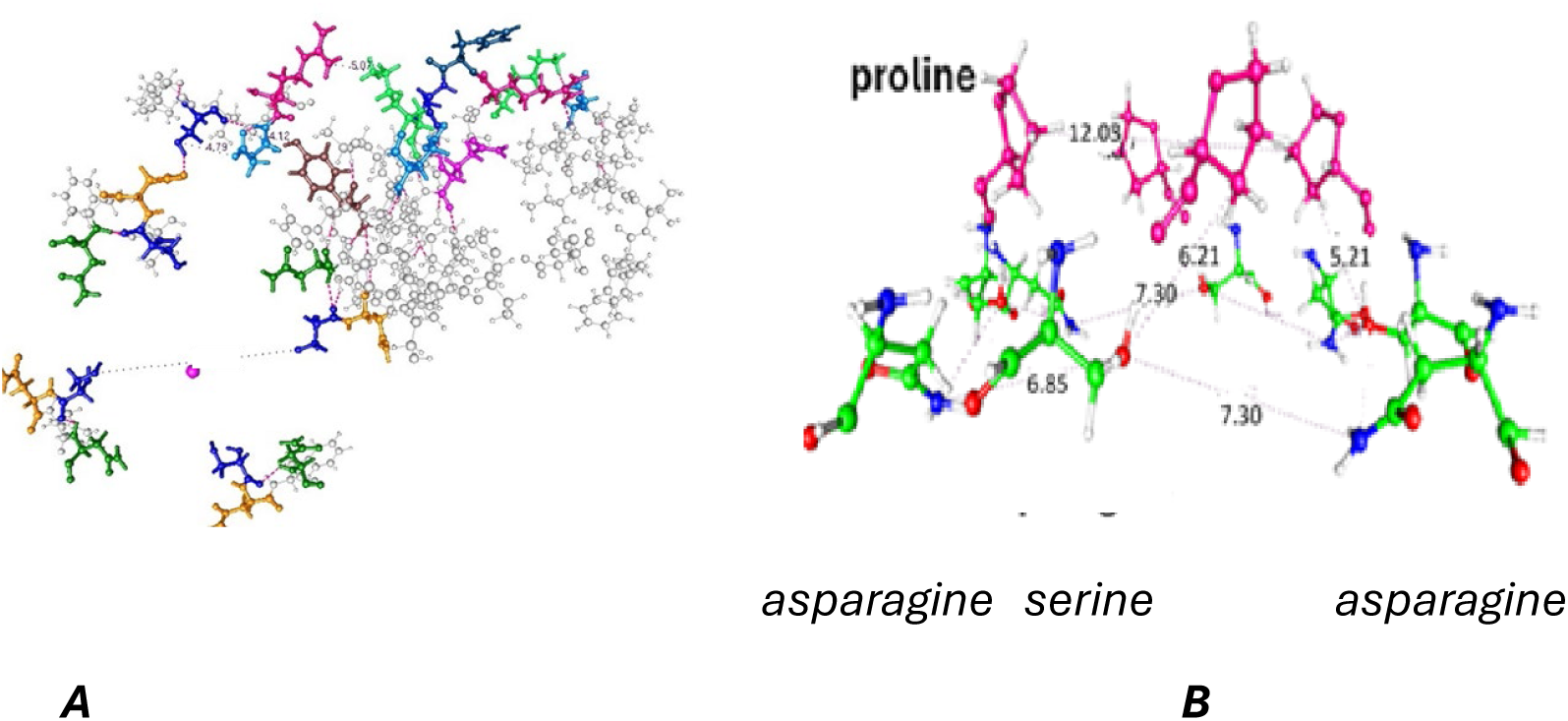
**A**) Looking at the channel structure from the intracellular end, looking up, with a purple potassium in the middle of the pore. The distance between the opposite serines facing the pore is 15.9 Å. The three amino acids at each corner of the figure are asn,ser,asn, with ser blue and the asparagines yellow and green). **B** Detail: Parts of two diagonally opposing domains of the four at the open gating region, seen from the side. The lower left has one of the asparagines, and the serine. Some key distances are shown. A K^+^ would proceed through the space from the bottom toward the upper left. The intracellular space is at the bottom here, the pore cavity at the top. As noted above, the distance across at the top is roughly 12 Å, at the bottom around 15 to 17 Å, depending on exactly how the distance is defined; these distances go to the outside of the figure, with internal distances (Å) along the dotted lines as shown. The taper fits the ion entering, hydrated, from the bottom, and leaving mostly dehydrated at the top. The vertical distance is roughly 6 Å. Water and ions are omitted in this figure. This is taken directly from the pdb file 7Sip, not optimized; the optimized distances are quite close to these values. The proline carbons are green, hydrogens white the water red (oxygen) and white (hydrogen), the K^+^ purple, the Cl^-^ red-brown, and nitrogen blue

Given the distances in Fig. 1, fitting fully hydrated K^+^ through the upper part of the gating region might be marginally possible at best, but almost certainly would face a very large energy barrier. Also, as the ion must lose at least four water molecules on passing through the channel overall (generally entering with 6 waters, it leaves with at most two), proceeding with more water molecules would imply a countercurrent of water for which there is not only no evidence, but which would require a major, and somewhat implausible, rebuilding of the gating region as each ion passes.

*Competitive Structures:* We need to introduce a new term, *competitive structure*, illustrated in Fig. 2.

**Fig 2:**
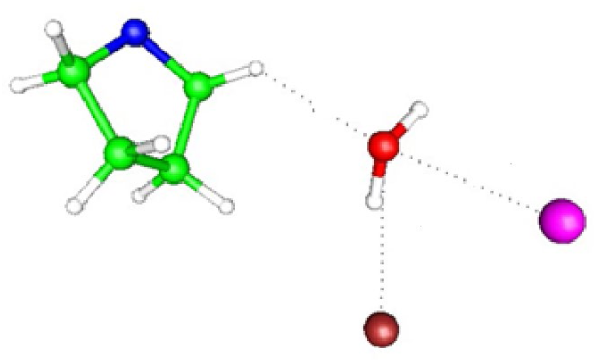
An example of a competitive structure: As in Fig. 1, the proline carbons are green, hydrogens white, the water oxygen red, the K^+^ 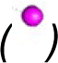 purple, the Cl^-^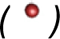 red-brown and nitrogen blue. Distances in this example: proline H to water O: 3:35 Å, O to K^+^ 2.97 Å, O to Cl^-^ 3.33 Å. Structures like this are found regularly in the computation; this case is cut from an actual optimization and the competition for the water molecule here is among the K^+^, the Cl^-^, and the proline. The electron density is determined between the relevant atom of the water, and the relevant one of the three other species.

We will refer to “competitive structures” from time to time, when needed to consider how the water molecules are stripped from the K^+^. This is central to the mechanism by which we get largely dehydrated K^+^ ions. The removal of water from the K^+^ ion depends on the interaction of the water with either another ion, or a side chain of the protein, or even another water molecule, or two waters.

Fig. 3 shows a competitive structure with two of the prolines showing.

**Fig. 3:**
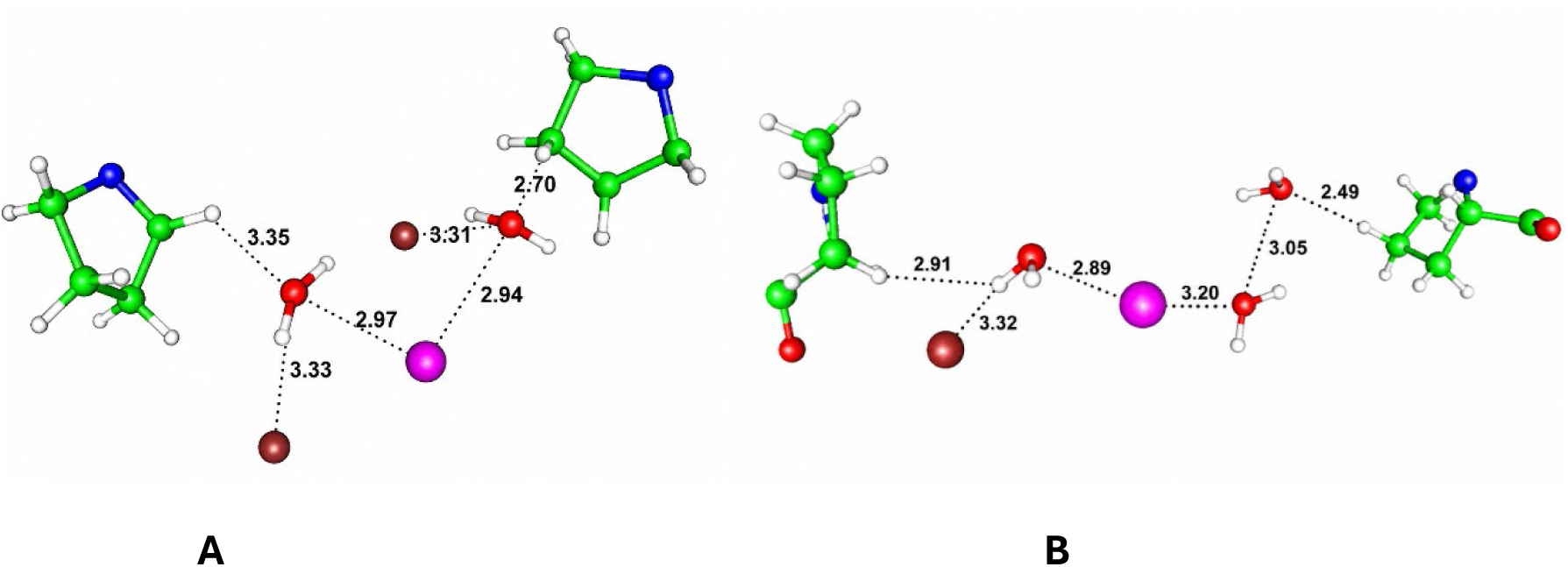
The competitive structure for two views of one case with two prolines showing, cut from the optimization of the wild type without protons, upper K^+^. Atom colors are as in previous figures. **A**: Here, the competing structure shows how the potassium ion can lose two waters from its hydration shell. **B** the same K^+^, but the figure is rotated and shows the arrangement along a different axis; the competing structure on the left removes one of the water molecules; on the right, we see the case in which the water remains with the K^+^ as the hydrating water’s neighbor is another water, not tightly held, so that the K^+^ keeps that one water.

A potassium ion enters the gate hydrated by six waters (occasionally one more or less; we will see cases with only five) and leaves the pore at the other side of the membrane with one, or at most two; this has been measured using streaming current for the bacterial KcsA channel and the K_V_1.2 channel; the ion is therefore hydrated with only one or two water molecules when it leaves the selectivity filter (12, 13). Therefore, it must be mostly dehydrated to pass through the pore, and this must happen at the gate, as there is no other location where the extra water can be stripped off, at least without creating a major countercurrent--even if the ion is partially rehydrated at the central pore above the gating region, an ion that arrives at the gate with six waters would require a countercurrent of four or five water molecules as the hydrating water is removed at the selectivity filter. The central pore can maintain a steady state if the number of water molecules accompanying an arriving ion equals the number with which it leaves to enter the selectivity filter, that is, one or two water molecules. This implies that the gate, when open, cannot be simply expanded to a large diameter that would let a hydrated ion through. Based on the calculations described here, the approximately 12 Å opening at the gate transition to the pore cavity must be adequate to accomplish this. Protons play a role in gating many channels, and we will see they play a major role here. The region between the asparagines or glutamines at the bottom of the gating region that are fairly well conserved (see Fig. 1), and the prolines above, is large enough that it could contain about ten water molecules at bulk water density, but can contain slightly more, together with ions, as a confined space with hydrogen bonding to the protein. Bulk water density is low because of the ice-like hydrogen bond network that holds molecules apart. There is a substantial literature on confined water (14–20). Some of this is not so relevant, as it concerns water that is confined by carbon nanotubes or graphene sheets. However, there is also considerable work on water confined near protein cavities, or with structure determined by water-boundary interaction (21–23). For example, a silica interface can be tuned to adjust free energy and affect a reaction (24); if a silica interface can be tuned, the much more complex protein interface can surely be tuned as well. With a relatively high density and several different species competing for bonding to water, the resulting structure of the confined space depends on the interactions of water and ions in the confined space. There is competition between K^+^ ions and protein and counterions (here, Cl^-^) for water molecules from the hydration shell of each ion, and the structure depends in large part on the outcome of the competition.

All this said, the results of the optimization were somewhat surprising. We had expected a simple effect, including a tight network of hydrogen bonds, together with a small electrostatic barrier to the movement of positive ions. This would be altered by the presence of protons, which could prevent the dehydration of ions while creating a substantial electrostatic barrier for cations. When protons are absent the results of the competition would be very different, and the ion would be able to move. The consequences of the rather high concentration of ions are discussed below; the concentration increases to a nearly saturated state at the gate. The optimizations produced a more complicated result than we anticipated.

There is a particularly interesting set of experiments from the Swartz group in which there is a mutation of the gate prolines (Fig. 1) to each of several amino acids, among them serine and aspartate (25, 26). Serine has a very small side chain, while aspartate’s is large. On the standard structural models, this should suggest that the serine mutation should be constitutively open, while aspartate should be mostly or completely closed. The opposite is found; the serine mutation is closed at all physiological membrane voltages, while the aspartate mutation is constitutively open. In this work, we present calculations on the aspartate and serine mutants, showing how aspartate opens the channel, and serine closes it. Our hypothesis suggests that cysteine, with a short side chain, should be mostly closed, while asparagine should be mostly open. This is in fact what is found experimentally. Because the gating picture that posits a mechanical blockage at the gate to close the gate would predict the opposite, a gating model other than the mechanical blockage of the pore opening at the gate is required. We could not do calculations for cys and asn for want of resources, but the experimental results fit well within what our hypothesis predicts.

This suggests that there is a set of considerations that have been omitted from the mechanical approach to channel gating. There is water in the gate region, as well as in the intracellular solution. The ion must pass through this region, losing most of its water of hydration. At the same time, it disturbs the hydrogen bond network, which must be able to break and form again for each passing ion; however, if there were a countercurrent of water, the required rebuilding of the gating region would be far more drastic than just a hydrogen bond rearrangement. There may be more than one K^+^ in the gate region and below. To avoid having too much charge in a small region, there are likely to be about as many counterions. These would be selected against in the gate region; several mechanisms are possible to contribute to this. It is generally difficult to tell where the water is in a cryoEM or X-ray structure, although a few positions may be sufficiently tightly held that they are present in all channels, and thus visible in the structures. However, while density for the oxygens may appear, the hydrogens will be invisible, so it will be difficult to tell water from a monovalent ion, and the orientation of the water dipoles cannot be determined from the structures. In calculations of hydrogen bonding this orientation is critical. Because the ions move through, the open structure must allow for dynamics; much of the water may be in different orientations, and probably positions, in an ensemble of channels. Structural water (at least the oxygen) may be visible, but the apparent electron density it provides may be interpreted as part of the protein. The same is true of the ions, including counterions. The ions are necessarily mobile in the open structure, and we have few closed structures determined from experiment. Therefore, it is likely that the structures will fail to show most water or ion positions. While there is little literature on the gate water structure in K_v_ channels, there is a fair amount in TRP channels, especially TRPV1, and work on these channels is consistent with these comments; TRPV1 in particular is gated by a ligand, capsaicin, by temperature, and by protons. The effects of protons have been studied by a number of groups. For example, Hazan and coworkers showed that TRPV1 requires 4 protons to gate (27). A number of groups have investigated the role of protons both in gating and in inhibiting gating in TRPV1 (28–32). In particular, protons stabilize the closed state of TRPV1 (30). We are primarily concerned here with K_v_ channels, but the domain structure of TRPV1 is somewhat similar to that of the K_v_ channels, in spite of the alternate gating modes involving temperature and capsaicin. These are not the only known cases in which protons couple to gating. For example, chloride (Clc) channels, as well as some transporters, show proton gating, mostly in the direction of protons opening the channel, opposite to what we find for the K_V_ channel here (33); protonation of glutamate appears to be key to the mechanism (34–36). While protons open some channels, there are conditions, and mutants, in which it does the opposite, closing the channel. The somewhat analogous structure is an additional argument for considering a possible role for protons in gating the K_v_ channels as well.

In earlier work, we showed that protons could reasonably be expected to move through the voltage sensing domain of Shaker, or of K_V_1 channels, and then to the pore with two possible paths through the linker(37) (38). In recent reviews, we have compared a proton gating model to standard gating models (11, 39).

A review by Zhang et al (40) considers the experimental properties of aqueous solutions in confined spaces. However, there is little in the way of directly relevant quantum calculations in the literature so far, although more is appearing. As always, MD calculations with classical potentials are not very useful for understanding what is happening at the gate, as the force fields are not calibrated for these confined conditions. Water rotation is (somewhat) hindered in confinement, and the dielectric constant is lower than 80—how much lower would require an extended calculation, and the necessary assumptions for this calculation are still somewhat uncertain. A low dielectric favors clustering of ions. However, there is enough work so far to show that we can reasonably expect clusters of ions in the confined solution at the gate, of ions with water, including clusters that are not well ordered. The density of these solutions may be slightly greater even than that of bulk solutions as the clusters bring the ions, often with water, together.

Water in nanometer sized protein cavities shows specific orientations, density, and hydrogen bond networks that depend on the way water is bound to the protein walls of the cavity, as well as how the water molecules either hydrate, or interact electrostatically, with the ions contained in the cavity, which are mobile. Being mobile, the ions require the networks to reform continually, although it is possible to calculate the network that corresponds to a particular static arrangement of ions. We are particularly concerned with energy minima in which a K^+^ is near the transition from gate to central pore, with another K^+^ not far behind, or just entering the gate from the intracellular solution, as these structures are where the difference between open and closed is likely to be clear. We have found the minima for cases that correspond to these requirements, and consider the implications for gating.

Water properties are complex. The longer range interactions, producing correlations among the waters (41), can also be seen in the confined water. In the literature, there is increasing attention to water in proteins; for example, one review uses rhodopsin as a test case (42), showing in particular the importance of water in channel-like systems. Protonation is also shown to be important there; in general terms, that review offers considerable support for the results we report. The more general properties of water have also been reviewed by Brini et al (43) who go through the complexity of the ways water can interact, not necessarily in a confined space. Water is still a nearly spherical molecule interacting through van der Waals and electrostatic forces, but unlike simpler molecules, its orientation is also critical—the dipolar forces are directional, and, more important, hydrogen bonding is specific as to orientation and critical in terms of electron density distribution and partial (non-covalent) bonding.

Because of the diminished dielectric response of the confined water, ion pairs that would not form in bulk, at least not in fairly dilute solution, such as H^+^ - Cl^-^, may exist in the confined space. Also, there are solvent separated ion-pairs to consider, with a water molecule separating the two ions that are paired; the results section shows some examples. Bulk water is 55.5 M, and there should therefore be about 28 water molecules per ion in a one molar solution of 1:1 electrolyte. However, the saturation concentration is nearly an order of magnitude greater, and the density of water itself can be greater as the walls of the confined space compete with the ions in the confined space, and with other water molecules, for hydrogen bonding to a water molecule. Deviations from the Debye-Huckel limiting law (the limit being infinite dilution) start at concentrations, for 1:1 electrolytes, above around 0.1 M, when there are about 280 molecules of water per ion. At concentrations much above this, the water cannot be treated as a continuum. In fact, the ionic strength of the intracellular solution is even greater (ionic strength includes all ions) than that of the KCl alone. The system must be considered with molecular bonding explicitly determined. When we talk about the local interactions, we do not mean that the surroundings are to be ignored, but that the local interactions must be treated differently than they would be in a bulk solution, especially if we are thinking of dilute solutions. At the gate, water molecules outnumber ions by much less than 280:1. In a saturated solution the ratio is more like 6:1, and at the gate we expect to be closer to the saturated solution than the dilute solution. The P475D mutation not only keeps the gate open but increases the conductivity by about six-fold over wild-type (44). Additional aspartates in the pore do not increase the conductivity any further.

Interactions with the ions may be largely electrostatic, but not entirely. For example, chloride ions in NaCl can form clusters (45–47) near the wall, approaching nucleation density (48). Together with the water, two ions with the same charge may be part of the same cluster, the electrostatic effects overcome by water molecules in these conditions. Electric fields can have competing effects, promoting nucleation of clusters but also driving ions apart (49). The fields, if large enough, can even lead to ionization via the Second Wien Effect (50). Competition between water-water and water-surface hydrogen bonds can lead to a more nearly two dimensional network, with higher density (51). New results also suggest that amide hydration in proteins differs from ordinary hydration (52). There are also changes in behavior with D_2_O. It is known that at least some sodium channels gate differently with D_2_O (53); other solvent effects on gating are also known, especially osmotic effects (54–57), Many other studies have been done in micelles or zeolites or other systems, but what these have in common with gates is limited to a demonstration of how the confined dimensions alter the properties of water. However, the fact that there is a solvent effect, and a D_2_O effect, is already enough to tell us that the water cannot be neglected; the fact that amides in proteins solvate differently tells us that the gating mechanism must involve not only the water but the relation of the water to the protein (58, 59). In the Results section we will see an example of the interaction of two K^+^ ions that appear to be bound through bridging water molecules.

So far we have not discussed quantum effects, but we can quickly see that these are not negligible; start by noting that the de Broglie wavelength of a proton with momentum corresponding to thermal kinetic energy at about 300 K is over 1 Å, in fact closer to 1.5 Å, thus comparable to the length of a hydrogen bond. The entire “diameter” of the open gate, at its narrowest, would allow for a pair of water molecules (four hydrogen bonds) and an ion. If the hydrogen bonds are delocalized to the extent that their positions are not well defined on an Angstrom scale, and water has electron clouds that are somewhat widely distributed, the entire gate can be a single network with a gel-like arrangement of extended electron clouds. There has been considerable previous work demonstrating such effects. Concerted hydrogen bond breaking by quantum tunneling, in a water cluster, the hexamer prism, has been described (60). Several groups have reported quantum delocalization in acid-base chemistry (61–63). Low temperature coherent structure with protons was demonstrated in a confined structure (14 Å) by Reiter et al (64), and ice-like arrangement in nanotubes, with structures that can be tuned by tube diameter, have been demonstrated (65). These examples suffice to show that the behavior of water in confined spaces needs to take into account quantum effects, and proton delocalization; added protons can reasonably be expected to significantly alter the structure of the water at the channel gate; this effect is apparent in the calculations reported here. The possible resonance effects in a network with cycles is mentioned briefly in the discussion section. However, the added protons are mostly tied up in hydronium ions, which may combine with water to form Zundel ions (this appears to be the case in several of the calculations), so that the properties of a free H^+^ may be less important.

It is therefore reasonable to expect an aqueous solution in the gate to have a fairly high density, ion pairing that would not exist in bulk solution, and a strong effect of added protons. The protons change the electric field, in addition to having a major effect on the hydrogen bonding network. They can also delocalize, creating a kind of linked network that must be “tied” (hydrogen bonded, or otherwise anchored) to the protein that constitutes the boundary of the confined space. Explicit quantum calculations are necessary to sort out the contact or solvent separated ion pairs, and the competition between the wall and the water for non-covalent bonding to the ions and the other water molecules. There is competition for which species may hold the water molecules that arrived hydrating a K^+^; the competitors include the protein wall, the K^+^ itself that may hold on to one or two of its original hydrating water molecules, the counterion, and other water molecules. The latter may pair with the proton, as well as other water, but not the cation; the water molecules may be part of a solvent-separated ion pair involving the counter-ion; the water in the solvent separated ion pair can be one of those hydrating the ion. Our new calculations show that the presence of protons specifically in the gating region can block the passage of the cation in the wild type channel, principally by affecting dehydration of the K^+^; with protons present, the ion is insufficiently dehydrated to pass through the upper part of the gating region and reach the pore cavity. This does not contradict the fact that protons are known to favor the open state of the gate in other channels, where they may affect the orientation of a side chain, or interact with a carboxylate side chain, among possible alternate effects. The mechanism by which the protons act here is quite different.

In earlier work, we considered the possibility that protons could constitute the gating current, rather than motion of the S4 transmembrane segment of the voltage sensing domain, suggesting a proton path from the S4 transmembrane segment through the S4-S5 linker (38, 66, 67). A new paper offers additional details as to the way this may happen (68). The VSD resembles the H_V_1 channel, so it is not surprising that it transmits a proton in response to a change in field, and it is not difficult to find a path between the VSD and the pore entrance; these were the subjects of previous papers. The chloride anion is on the border between chaotropic and kosmotropic (i.e., structure-breaking vs structure-making). Methane sulfonate, which has been used in some experiments, is strongly kosmotropic, and the anion of a strong acid. It would be interesting to see what happens with a chaotropic anion, but the experiment would be complicated. Also, the anion of a weak acid might hydrolyze, producing an additional unwanted complication. At present therefore we stick to chloride in these calculations. Local interactions, including the electron density differences between ions, counterions, protons, and water, allow the dehydration of the ions. We can use electron density differences to determine whether a given water molecule that may be around 3 Å from a K^+^ is actually hydrating the ion, or whether it would stay behind if the ion moves ahead in the membrane, more or less bound to another species. It is here that protons play a key role. While we concentrate on these local effects in single channels, we are not arguing that the confining spaces do not also play a role in the overall behavior of groups of channels; there are also effects of lipids (69). However, gating cannot be understood if we do not examine in detail the interactions among ions, water, and the confining boundaries in individual channels. Further support for the importance of dehydration in moving ions through nanometer pores has been provided by Wang and Sun (70); they found that electrically driven transport is also dependent on the diameter of a hydrophobic tube through which ions are transported.

When additional protons are present, there is a major rearrangement of the entire WT gate aqueous configuration, and we see how the K^+^ is held up by failure to dehydrate, when coupled to the water and H_3_O^+^, through an altered network of ion pairs and hydrogen bonds; the interactions with the counterions change. When the channel is open, the current flickers; the exact reason is not so clear, but it does suggest that the open state exists on the edge of transition to some sort of closed state, possibly only a transient state. We note that there are a number of channels with conductivity affected, up or down, by protons in various positions on the channel protein (71).

There is some additional evidence in the literature that is relevant to the hypothesis that we present here. Low pH has been shown to close the K_v_1.2 channel, which strongly suggests that protons do (72), and this has been supported by simulations (73), although the simulations suggested dewetting the cavity to close the channel with added protons. There are also channels other than K_V_1.2 or Shaker in which S4 does not move; one of the clearest instances is a sodium channel (74). Another case in which protons are involved in closing a potassium channel is in a cytochrome (75). The Zhang et al Eag2 channel result, cited above, is another example (6). Therefore, multiple instances are known in which protons can gate a channel, or that S4 does not move as part of gating.

### Summary

It is clear, from the fact that the ion enters the gate with 6 ±1 hydrating waters, and leaves on the extracellular side with at most 2 (12), that the K^+^ ions must be partially, indeed largely, dehydrated. If the ion passed through the gate fully hydrated, and thus entered the pore cavity, there would have to be a counter current of water; not only is there no evidence for such a thing, but it is clear that it would have consequences incompatible with observations. The mechanism by which dehydration happens must be worked out. We hypothesize a key role for protons, with calculations showing how this may be determined as well. Lipids, or clustering of channels, may affect the transport of protons from VSD to gate, or have an effect through some other mechanism, but the question of how the ion must interact locally in the gating pore is central to understanding gating. It is this question that we address here.

We can therefore restate the hypothesis.

### The proton gating/ion dehydration hypothesis

The hypothesis has two parts: *First*, the K^+^ must be partially dehydrated; this is almost forced by the fact that the ion arrives at the gate hydrated with, normally, six water molecules, and the ion leaves the channel with at most two water molecules. Four water molecules (sometimes five) must be removed at the gate; if this happened further up the pore, there would be a counter-current of water, which seems wildly unlikely; certainly there is no evidence to suggest that such a thing exists, or could exist. Such a counter current would have to remove ions from the upper part of the gate region, and require that a conformation that included the necessary ion(s) in the required form for transition upward be re-established, which would be highly inefficient. It is clearly necessary to remove the water before the ion enters the water filled cavity in the center of the pore. The question is how this water is removed. The hypothesis states that in K_V_1.2 and *Shaker* (and other channels with the conserved proline-valine ring at the gate) competition for the water by counterions and the channel wall removes water molecules except for, at most, one above and one below the K^+^. A fifth water can sometimes attach to another species at the gate, so the ion might proceed with only one water molecule. The calculation can test whether the water molecules are stripped off or not.

The second part of the hypothesis concerns the gating itself. To close the gate, the hypothesis proposes that protons constitute the gating current; the positive charges that move from the VSD are protons, and not arginines. Some protons are trapped along the way, and four protons arrive in the gate although there is a gating current of ten or twelve charges. In other words, some of the protons remain behind in the path from VSD to the gate; not all the protons need to traverse the entire distance to the gate. With the protons, counter ions that participate in the dehydration of the K^+^ are partially neutralized, or ion-paired, so the water is not stripped away; in a sense, with the counterions largely prevented from participating in dehydration, the K^+^ “wins” the competition to hold the water, and therefore cannot progress into the pore. The difference must be subtle, as the current in most open channels, including the ones concerning us here, tends to be intermittent, so the channel, even when open, appears to be on the threshold of closing. The hypothesis explains an interesting experimental result. The conductivity of P475 mutants has been studied by the Swartz group (25, 26). The size of the gate opening is a significant part of what determines whether the ion is dehydrated or not. Because the properties of confined water are crucial, the exact conditions of confinement make the ion transit path different. The mutation P475D leaves a smaller opening than P475S. and is thus able to strip water with or without protons in the gating region, leading to a constitutively open (at physiological potentials) channel; the aspartate carboxylates assist in this regard. The mutation P475S, which leads to a geometrically more open gating region, in fact produces a channel which is non-conducting at all physiologically relevant potentials; this channel shows no competition for water molecules, so they stay with the K^+^, preventing conduction. In the serine mutant, there is sufficient volume in the gating region that the hydrated K^+^ can hold the water molecules, and thus fail to advance to the pore cavity. We have presented preliminary versions of this hypothesis earlier (11, 38, 66, 76, 77), and here provide much more specific evidence with new calculations, and with defined ion-ion, ion-counterion, ion-water, and ion-wall interactions, including electron density determination for these interactions.

We therefore postulate the following: 1) Dehydration of the K^+^ is required to traverse the pore. 2) The P→D mutant is constitutively open. The P→S mutant is closed at all relevant potentials. By the hypothesis, this is a consequence of the failure to dehydrate the ion. 3) The protonated wild type is closed, the unprotonated channel, open, and the difference is subtle, with the transitions near a threshold; the counterions must be included to understand the effect. 4) The counterions play a key role in the dehydration. Competition among the ions for water for their hydration shells is central to the dehydration mechanism.

### Summary of hypothesis

There are two main points: 1) The K^+^ ion must be mostly dehydrated to pass through the boundary between the gate region and the pore cavity 2) In the wild type channel protons prevent the dehydration of the ion, thus closing the channel

*The calculations test these points*.

## Methods

Quantum calculations were carried out on a section of the protein that includes the gate. For the wild type gate, 592 atoms were included for the cases without protons, and 593 atoms with four added protons; one water was removed. Calculations on two cases of each were carried out, one in which the initial positions of the K^+^ ions were closer together than in the other. Local minima are found by the optimization, and separate local minima were found for the two initial positions for both the protonated and unprotonated forms. Two mutants, P475D and P475S, were also optimized. There were again either zero or four protons for the P475D case; for P475S, even the zero proton case is expected to be closed, and the expectation was confirmed, so there was only one run, with zero protons. The 7sip pdb structure gives the starting coordinates for the open case, with waters and ions added, while the hypothetical closed case has four additional protons. Chloride counterions are added to balance charge; the open case is calculated with charge −2, the open case with four additional protons is calculated with charge +2. The volume calculated uses all four domains in a small region at the gate. The protein wall has the local amino acids (see figure 1) plus the added water, potassium ions, and chloride ions.

For the mutant cases, we included for aspartate 960 atoms, or 964 with added protons, for serine 961 (only with no protons) were included; the calculation was extended below the range considered in the wild type cases, with the addition of 70 water molecules, as well as additional ions, plus a slight extension of the protein. The wild type cases had only the two K^+^ that were in the gating region, with very little below it included. P475S was optimized with 2 K^+^ but P475D had 6 K^+^; as there are always 4 Cl^-^ the added K^+^ with aspartate kept the charge the same. However, again only two K^+^ entered the gating region, so the additions made essentially no difference, while taking up considerable computer resources. These calculations however served as a check that adding more to the bottom of the system was not important.

In each case, the optimizations were carried out using Gaussian16 (78) at HF/6-31G** level, with protein backbone atoms fixed and the side chains, the water, and the ions optimized. The optimized structures were then used in a single point Natural Bond Orbital calculation at B3LYP/6-31G** level, to obtain electron density for relevant bonds. The DFT method was too slow to be practical for optimizations.

### Limitations of the calculation

While the results of these calculations can be expected to be better than those of any classical simulation, (reasons for distrusting classical calculations were included in a review (11)), these calculations also have limitations as to quantitative accuracy. Given the available computer resources, one must make compromises in the way the calculation is done. While more sophisticated methods exist (79, 80), they cannot be used here because of the computer resources that would be required. Here, it was necessary to choose among the following: number of atoms included in the calculation, possibly the most serious limitation, the method (HF or DFT; if DFT, which functional?), and the basis set. Also, the protons are treated using the Born-Oppenheim approximation, and energy minimization means the calculation is valid, at least in principle, at 0 K, and we can reasonably expect the structure to be essentially correct there although done at HF/6-31G** level. The 0 K result should allow comparison to cryoEM and X-ray structures, both of which are done below the crystallization temperature of high-density amorphous water. The compromise chosen uses HF for structure, which is almost certainly good enough to give an accurate structure, but B3LYP/6-31G** for quantitative values in single point calculations The electron density values from B3LYP/6-31G** will be in reasonable relation to the correct values (81).

The NBO electron density values are taken as an approximate measure of the strength of the non-covalent interactions between atom pairs. There is a vast literature on non-covalent interactions, including hydrogen bonds. We do not review this literature here, but the interpretation of the electron density as approximately proportional to the strength of the interactions appears to be reasonable. We only use negative values of electron density as an indication of non-interaction, without quantitative significance. Because our conclusions only require approximate proportionality of the calculated electron density to interaction strength, the level of quantitative accuracy is less important.

## Results

The results of the calculations are shown in the Tables and Figs. 4, 5 and 6. The key figures are divided into two parts: much of the calculated region is included in one part, together with a detailed figure in which the key interactions are indicated; these are hard to see when most of the atoms are all shown together. The detailed figures show the kind of competitive structure defined in the introduction. We can see clearly the step in the dehydration, in which the water in effect is the prize in a competition between ion, counterion, sometimes another water molecule, and to some extent the protein wall. In one case, a hydrating water molecule also forms part of a cluster with three other water molecules, and would thus be removed from the hydration shell. In the WT no proton case, by hypothesis open, the chloride “wins” (*i.e.,* keeps the water) in enough cases to allow the ion to proceed with only two water molecules. NBO calculations at B3LYP/6-31G** level on the optimized structures showed the charge density overlap, thus accounting for the stronger interaction of some water molecules with the counterion, while the protonated case is more complex; see Tables 1 and 2 for wild type. In contrast, in the P475S case, at least four, very likely five, remain behind with the ion, effectively immobilizing it, whereas in the P475D case, the ion keeps only two, or just one; the electron density determines which is which. The electron density is of the order of 0.1 for the partial bond to water; in other words, this is a weak non-covalent bond. We will consider the implications of this below. With protons added, the last step for the proton in reaching the gate goes by way of the asparagines at the corners of the entrance to the gating region (Fig. 1B).

**Fig. 4:**
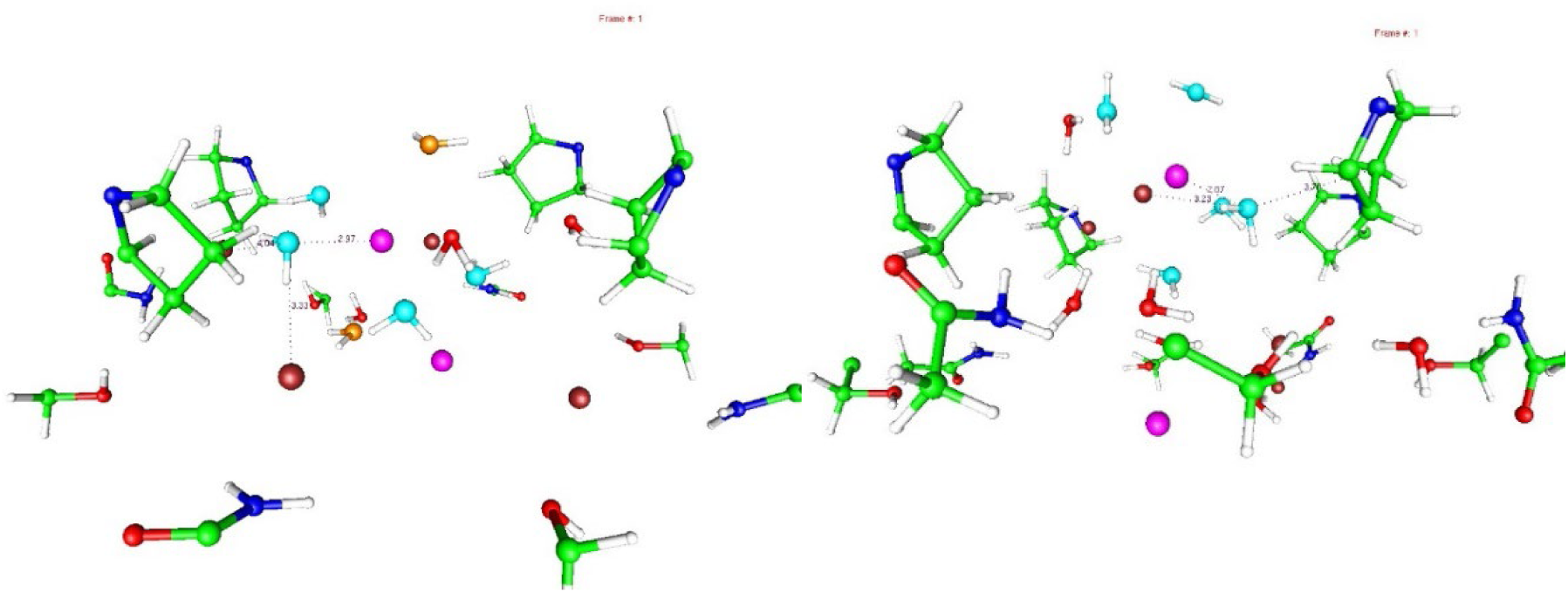
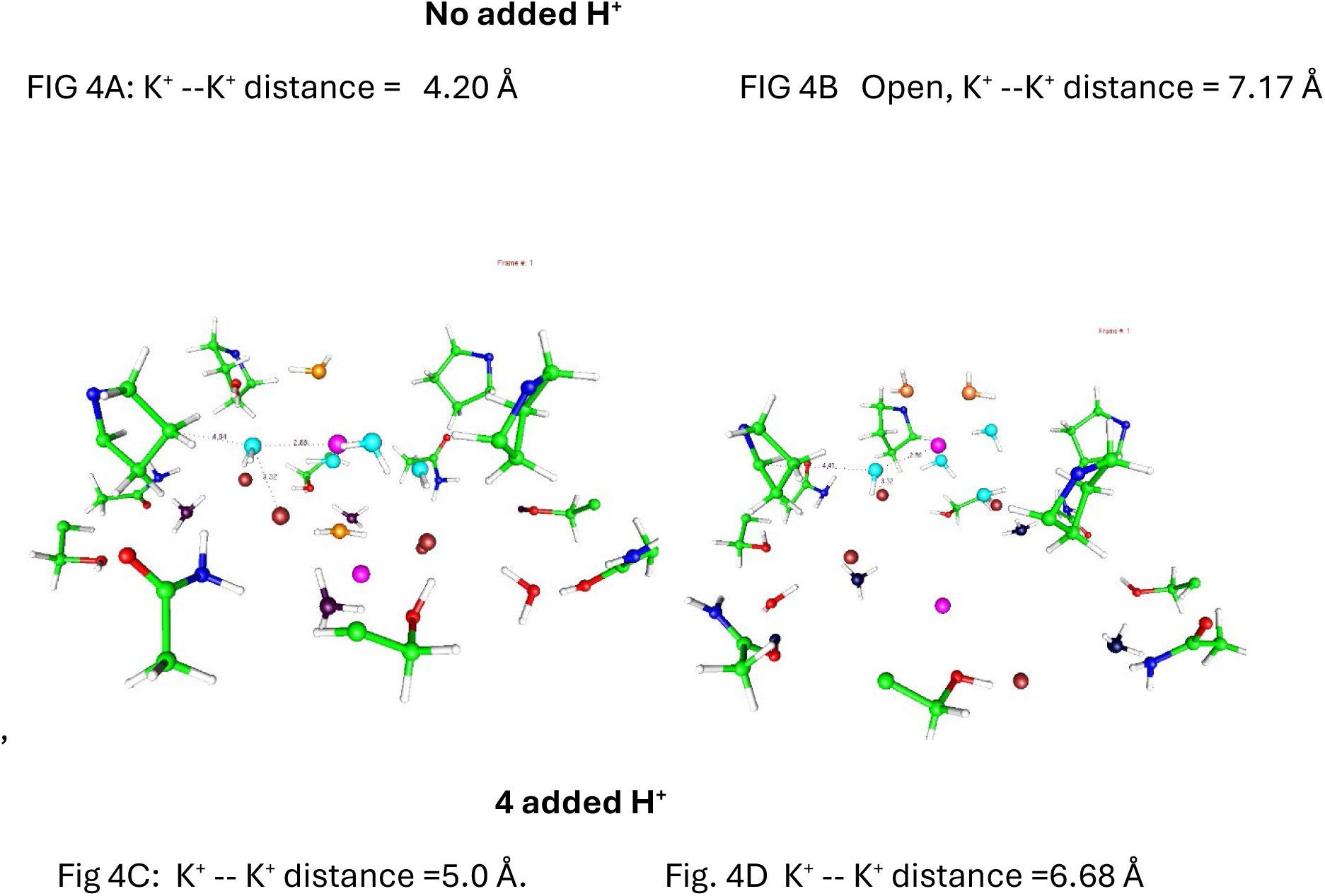
Wild type cases: The structure of the water network is very different when protons are present. In fact even starting the potassiums in different positions leads to a significant difference in the network in the two different minima that are found with protons, comparing Fig. 4C to Fig. 4D. Without protons, comparing Fig. 4A to Fig. 4B, the minima are more similar, and the K^+^ --K^+^ distances not so different; both show the kind of competing structures defined in Figs. 2 and 3. In all of Fig 4, colors are as before (K^+^ are purple, Cl^-^ are red-brown, C green, N dark blue, H white circles, most O, red), with one exception: water molecules that are directly involved with hydrating K^+^ or else are being competed off, are labeled separately. The light blue oxygens 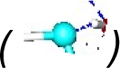 belong to water that remains hydrating K^+^, and are in roughly the same plane orthogonal to the vertical axis, as the upper K^+^. Water with orange oxygens 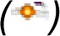 above and below the K^+^ are the part of the K^+^ hydration shell that is not removed, and may proceed (4A,4B, unprotonated cases) to the pore cavity. In the protonated cases (4C,4D), the oxygens of the H_3_O^+^ ions are very dark 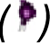.

**Fig. 5.**
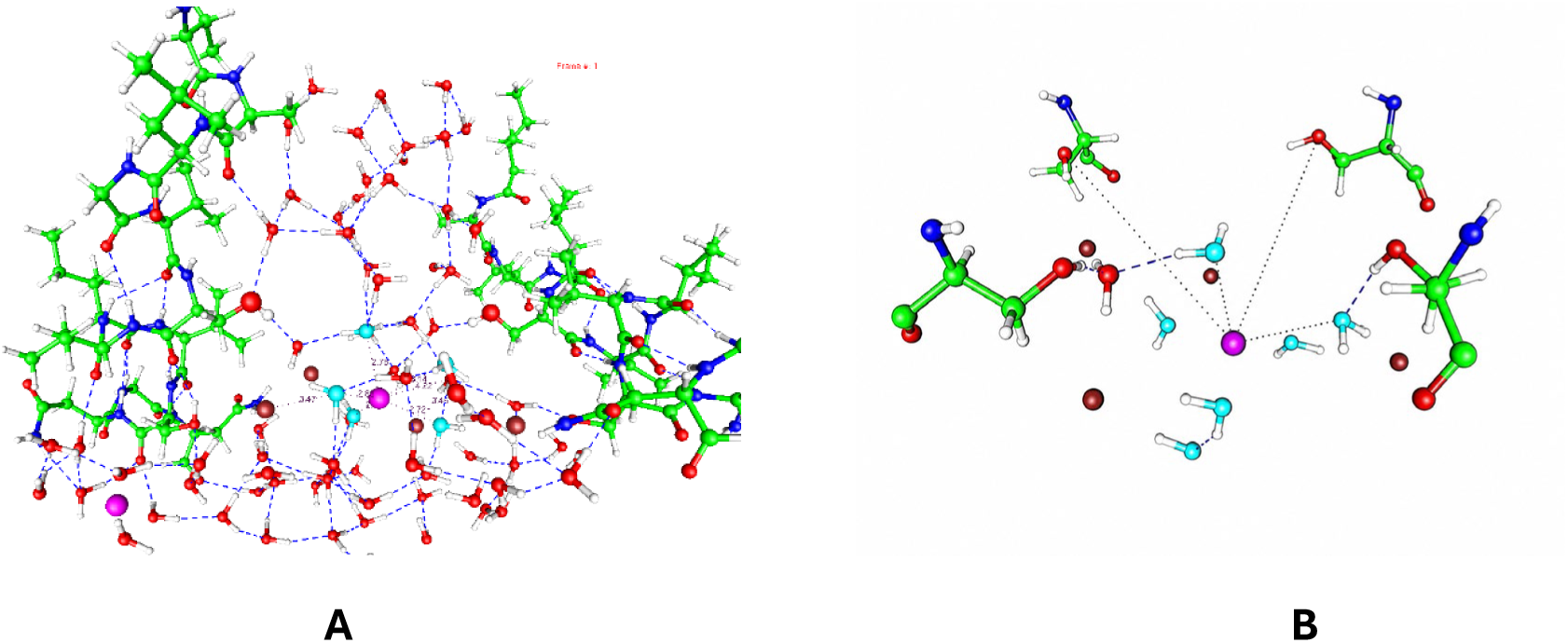
**A**: The serine mutant (2 K^+^): A section of the serine mutation cut from the central portion of the converged structure, with the light blue oxygens showing the position of the water shell hydrating the uppermost K^+^ ion (magenta). Four chlorines are reddish-brown 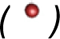. Examination of the positions of the water shell of the potassium ion on the right shows that only one of them is within hydrogen bonding distance of a serine -OH, and none is close enough to a Cl^-^for that ion to compete for the water either. Fig. 5B: A section cut from Fig. 5A, and reoriented slightly, in which it is possible to see that the potassium ion is hydrated by water molecules that are not close to any species (Cl^-^, -OH, or anything else) that can compete for the hydration water. Absent a competing structure, the K^+^ remains hydrated. When the distances and electron densities are examined, five of the waters remain with the ion, and one is removed by a competing water with electron density showing that that one water is removed; a second is a little less certain. Therefore, the ion cannot move from the gating region into the pore cavity.

**Fig. 6:**
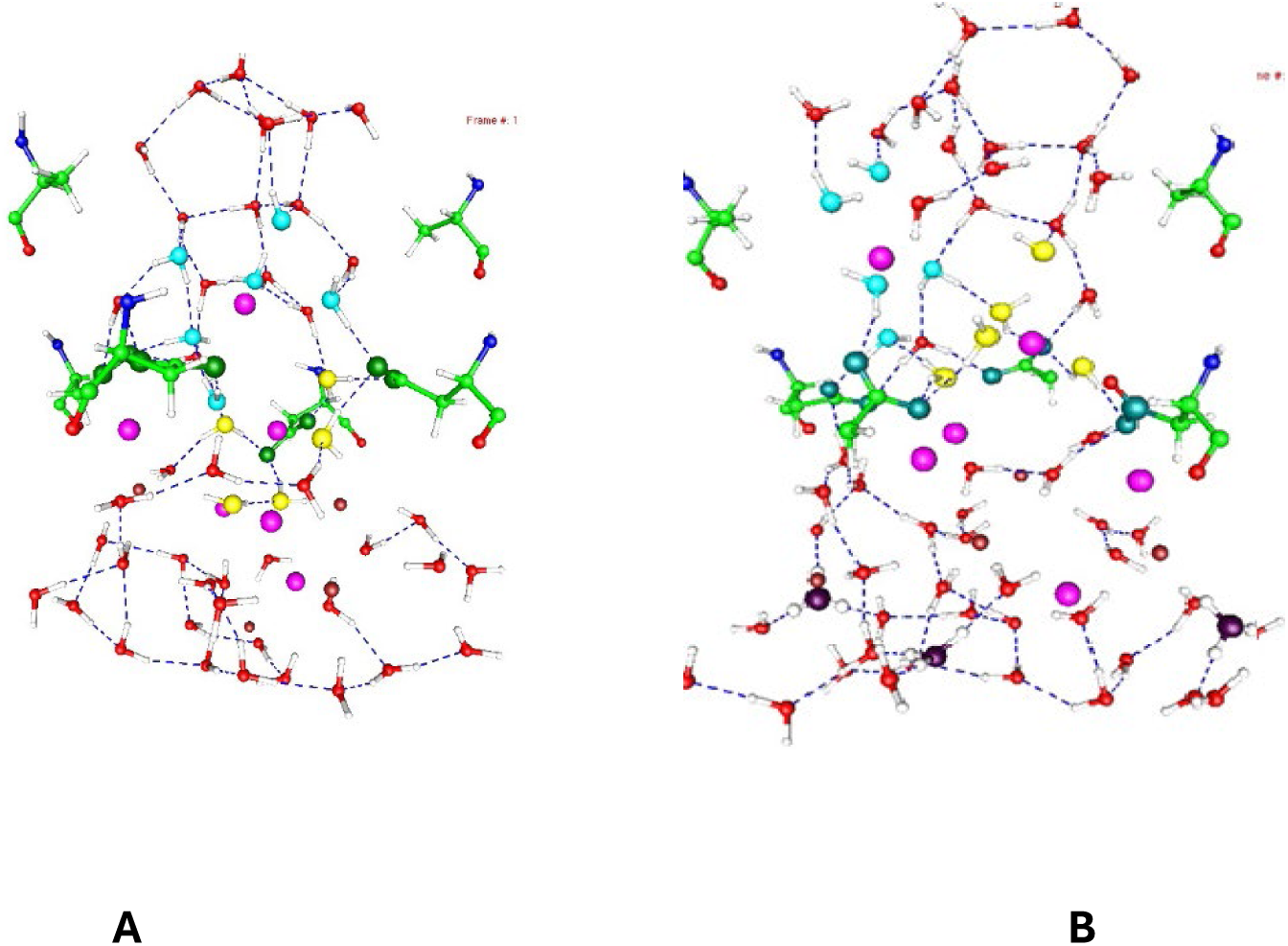
The aspartate mutation without (A) and with (B) four protons. This shows how the K^+^ ion passes through the gating region, derived from the optimization of the aspartate without and with four protons. Fig. 6A has 240 atoms of the 960 total in the calculation, figure 6B, 261. This allows the main part of the system to be understood. A middle shade of blue-green 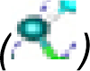 is used for the oxygens of the carboxylate groups. Light blue 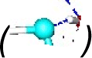 is used for the water oxygens that started as hydrating the upper K^+^ 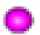 The water hydrating the lower K^+^ has yellow oxygens 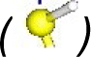. In Fig 6B, the proton’s O (i.e.,H_3_O^+^) are dark 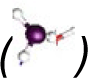. Only one Cl^-^ is visible here, as the carboxylates keep the Cl^-^ below the gating region. The other atoms have the same colors as in the earlier figures (C, green, other water: red and white. For the upper K^+^ the distances and electron densities are in Table 4: distances cannot be shown in the figure as they would be too crowded to read). In contrasting the appearance of the water network with and without oxygens, we see that the protons make a considerable difference.

**TABLE 1.**
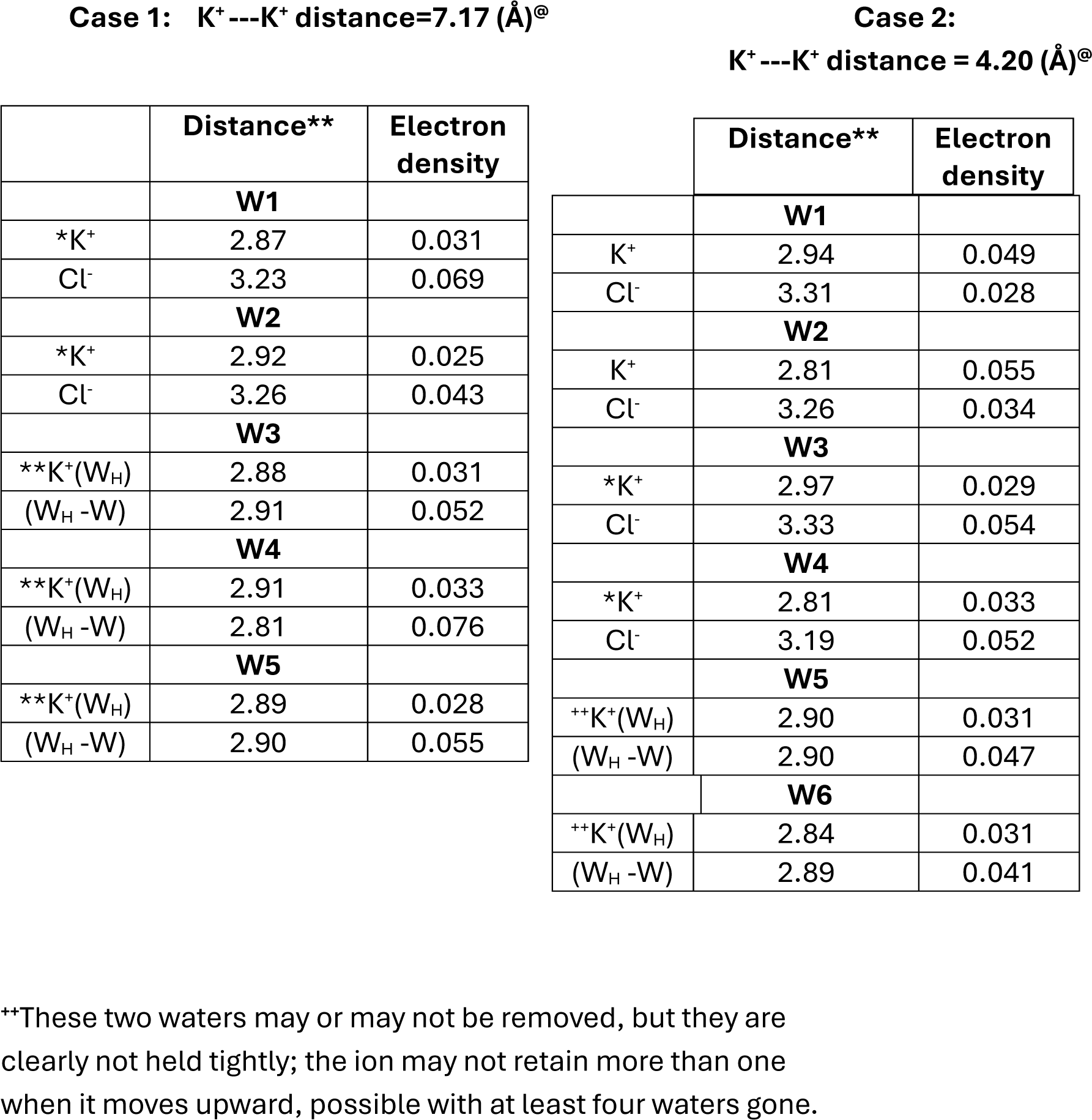
WILD TYPE; NO PROTONS.

**TABLE 2.**
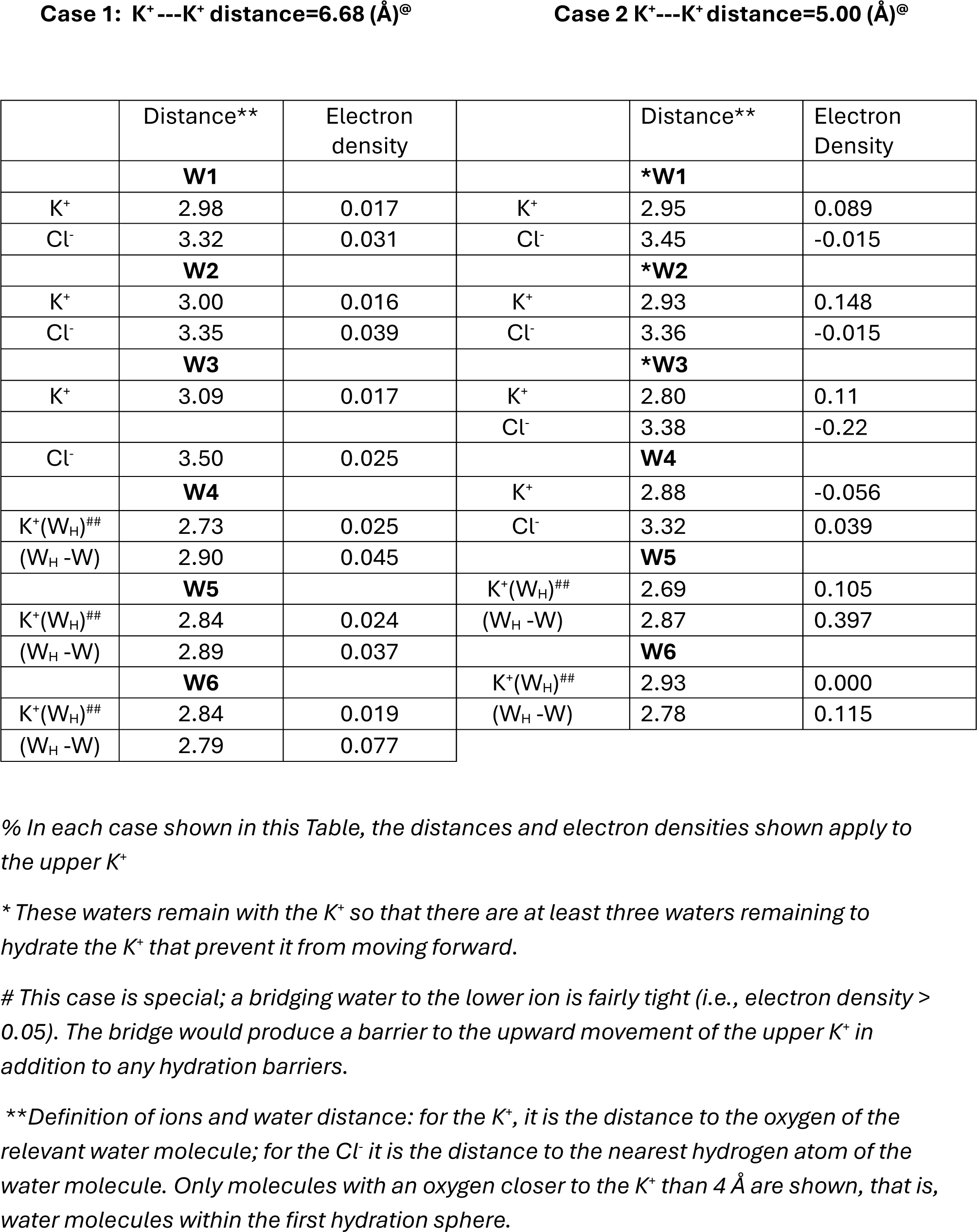
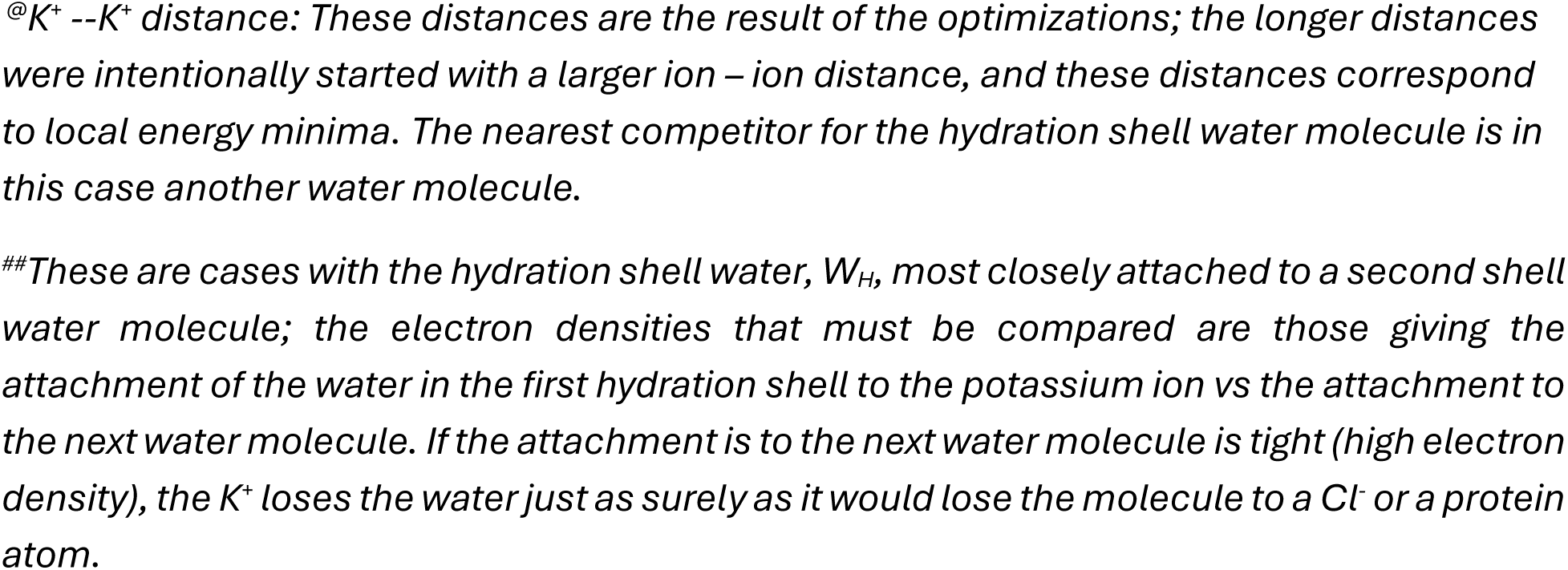
WILD TYPE: FOUR PROTONS.

### 1: Wild type

There is a tremendous difference between the unprotonated and protonated channels for the short K^+^ - K^+^ distances (4.20 Å, 5.00Å). Both with and without protons, the K^+^ ions are actually part of a single complex with two bridging water molecules. Two more marginal cases exist, although not identified explicitly as a water molecule bridging two ions. This accounts for the fact that the ions can be as close as they are.

For the large K^+^ - K^+^ distance (7.17 Å) with no protons, only two water molecules remain associated with the K^+^ ion. The hypothesis is that K^+^ needs to be dehydrated down to at most two molecules of water, and this does happen when there is no other close K^+^, consistent with the channel being open. In this case, only 5 water molecules constitute the first hydration shell, with all others too distant. When the two K^+^ are close (4.20 Å), two water molecules (shown as W1 and W2 in Table 2) remain hydrating the ion, and four are removed; for two of these, the competing species is another water molecule. When the K^+^ ions are distant from each other, and thus effectively independent, there is again a considerable difference. In both cases a single water bridges the ions, but it is weakly bound in the no protons case—the K^+^ -K^+^ distance is 0.49 Å longer when there are no protons present—and there are only two water molecules left in this case too. At this point the upper ion is free to move into the channel pore. With the protons present, the upper ion is still part of a network, and has three water molecules in its first shell. The protonated case appears to leave this ion immobile, unlike the corresponding no protons case.

However, if the water originally in the hydration shell is more tightly bonded to the outer shell water molecule, it will still allow the ion to proceed without the hydrating water molecule. The results, therefore, are in accord with this part of the hypothesis, which requires at least partial dehydration for the ion to move forward.

For the no proton case, where only two water molecules are within the first shell, the second shell begins past 5 Å distant from the ion (data not shown). A possibility that is not considered in these calculations is that there are more than four protons (the gating current has ten or twelve charges). In this calculation we assume that only four make it into the gate, with the rest trapped along the way, for example in T1 or the linker. If, say, eight protons made it into the gate, presumably the differences with the no proton case would be even more clear. If four protons suffice to close the gate, we assume that more would close it as well. More than four protons would require chlorides to balance the charge, and the consequent density (water plus all ions) in the gate would be high—the effective concentration would be higher than saturation would allow in bulk. It is of course possible that there are something between 4 and 8, perhaps 5 or 6 protons, at the gate. Given the four-fold approximate symmetry of the channel, we calculated the case of four H^+^, which proved adequate for the gating mechanism we are hypothesizing.

It is abundantly clear that, irrespective of the gating properties of the protons, they make a tremendous difference in the structure, or arrangement, of the water molecules in the gate region. That this would be irrelevant to gating seems highly improbable; gating involves passage of the ion through the network of hydrogen bonded water at the gate, a network that must rearrange as the ion passes through, and changes drastically when protons are added. It appears that the only way protons could fail to be involved in gating is if there are no protons that reach the gating region. This is a question we dealt with in previous publications, as discussed in the introduction, where we quoted our earlier work showing that protons would constitute gating current and can transfer from the VSD to the gate(37) (38).

The one set of data seemingly inconsistent with the second part of the hypothesis is the protonated case with the ions separated by 6.68 Å, the larger separation. Here the upper K^+^ appears to be weakly dehydrated, and none of the water molecules stick strongly to the K^+^, which therefore appears to be free to move in spite of the protons. It does appear that one water molecule bridges the two K^+^ ions (6.68 Å is very close to the correct distance for this, so this is not surprising), so that the upper ion would be tied down, unable to move forward. However, this is not conclusive evidence that the hydration of the ion is preventing it from moving forward. This may be the one case in which having additional protons might be relevant. We will consider this case below and in the discussion section; it seems anomalous, but not necessarily a serious problem for the hypothesis.

### 2: Mutations

The difference when the proline is mutated to serine, producing a channel closed at all physiological potentials, or to aspartate, producing a channel that is practically constitutively open, is shown in the corresponding figures and tables below. As noted in the Methods section, 960 atoms, or 964 with added protons, were optimized for aspartate; the gate was extended below the range considered in the wild type cases, with the addition of 70 water molecules, as well as additional ions, plus a slight extension of the protein. For the P475S mutation, which is closed at all potentials in the physiological range, only one calculation was done, without protons, and as Fig. 5 and Table 3 show, the ion is not dehydrated, with only one (possibly a second too) of six water molecules removed. The opening from serine hydroxyl oxygen to oxygen, diagonally across the gate, is approximately 11.4 Å, roughly 2.7Å greater than the carboxylate oxygen distances in the aspartate mutant.

**TABLE 3.**
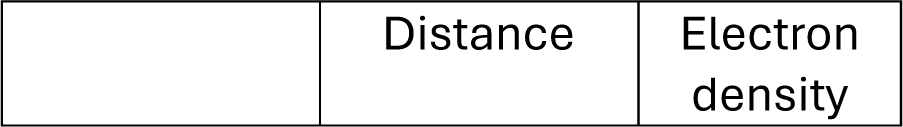
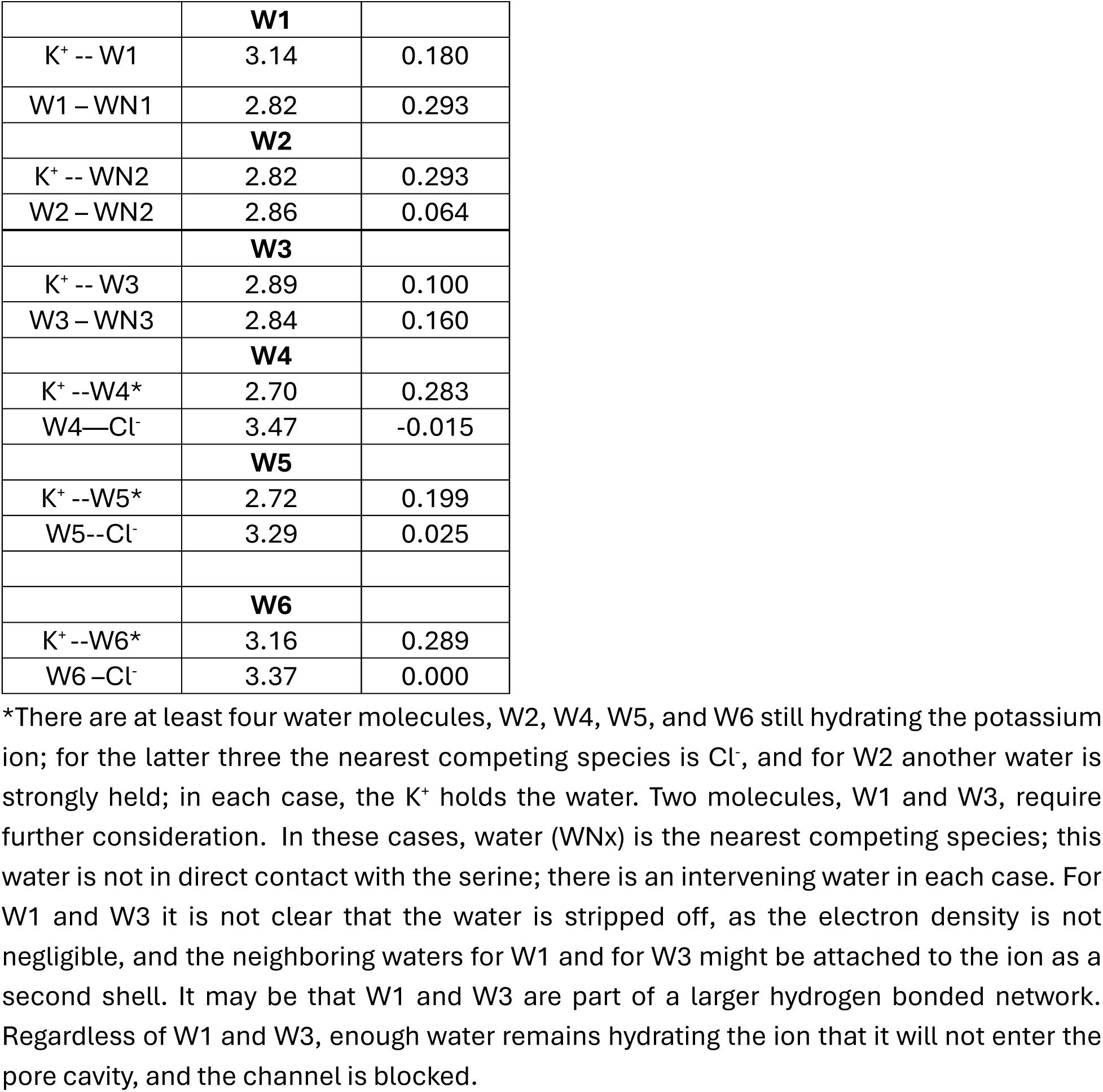
THE SERINE MUTATION, NO ADDED PROTONS Upper K^+^ ion with 6 nearest neighbor water molecules Nearest neighbor water with nearest competing species.

The added protons case with P475S has not been run because of the cost in computer resources, combined with the fact that the no protons case is already closed, so further closing it was not a high priority. In principle, protons might have a reverse effect and open the channel, but the probability seems small enough that it justifies being omitted; there does not seem to be any way that this could happen.

#### Aspartate

The results for the aspartate mutation are in Table 4 and the accompanying Figure 6.

**TABLE 4.**
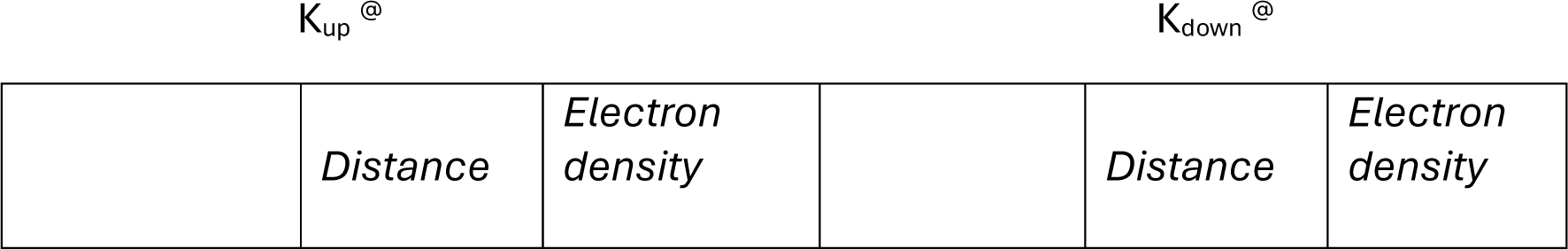
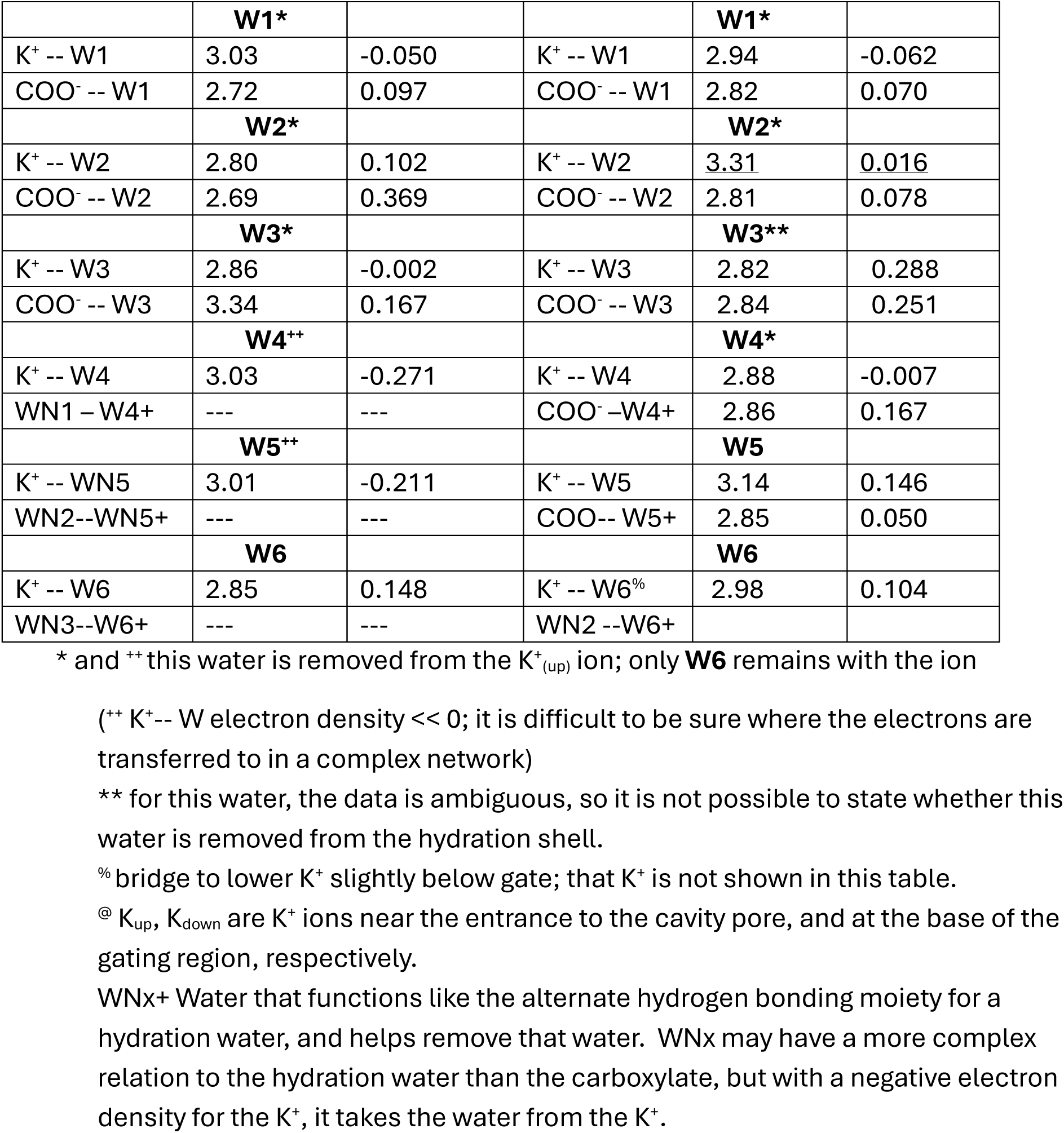
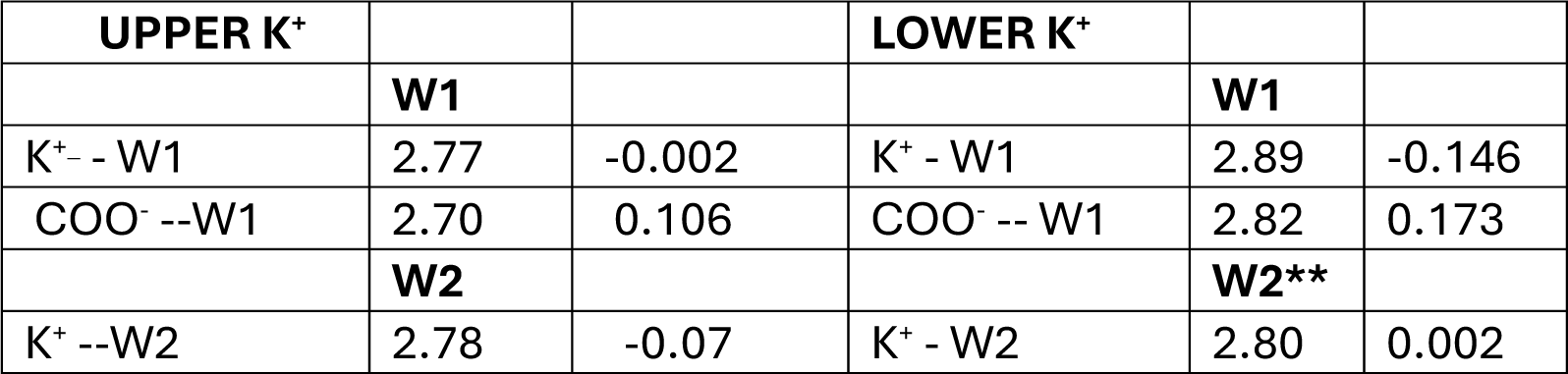
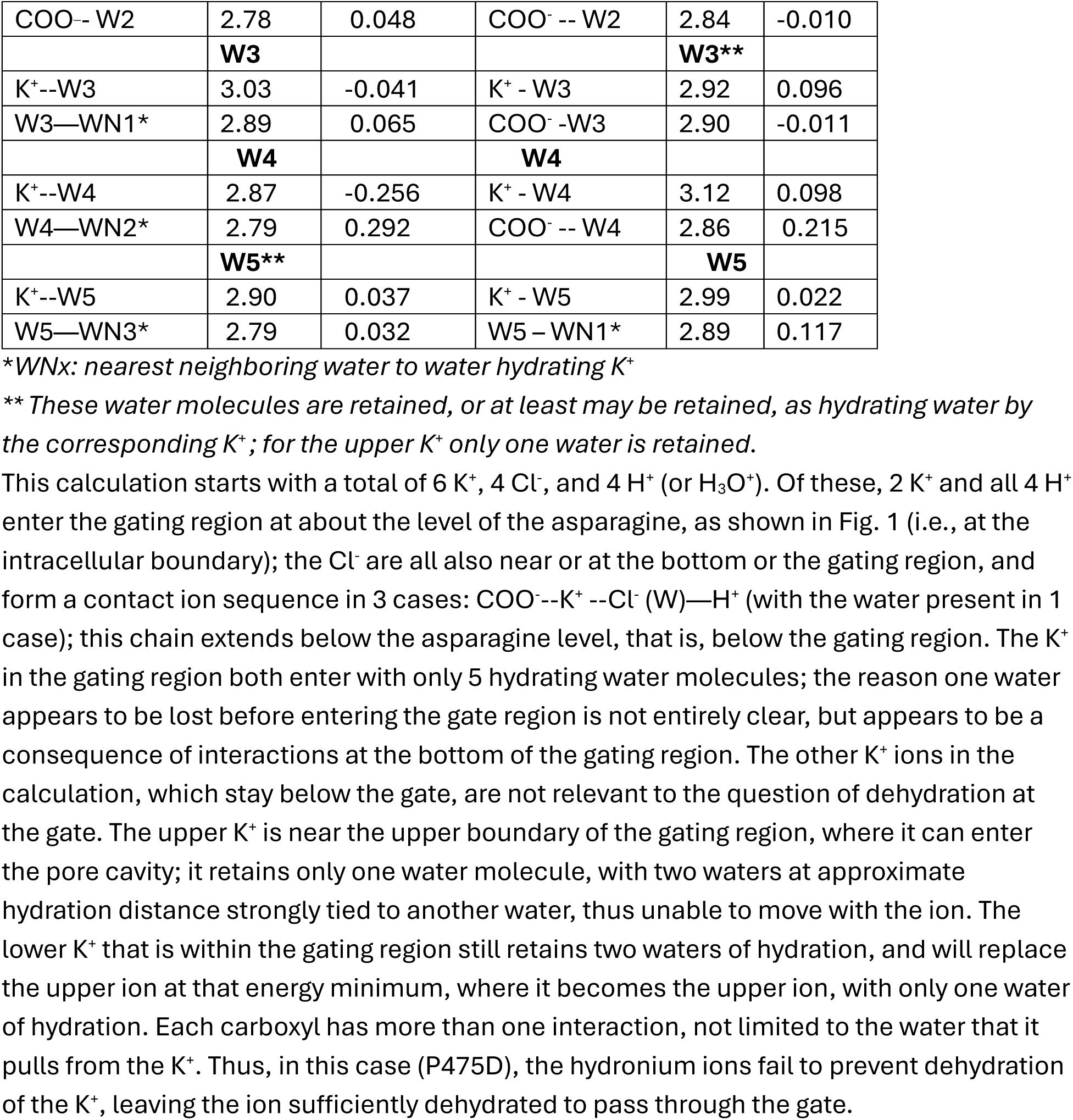
A ASPARTATE MUTANT, NO PROTONS ADDED, B ASPARTATE MUTANT, FOUR PROTONS ADDED.

This calculation starts with a total of 6 K^+^, 4 Cl^-^, and 4 H^+^ (or H_3_O^+^). Of these, 2 K^+^ and all 4 H^+^enter the gating region at about the level of the asparagine, as shown in Fig. 1 (i.e., at the intracellular boundary); the Cl^-^ are all also near or at the bottom or the gating region, and form a contact ion sequence in 3 cases: COO^-^--K^+^ --Cl^-^ (W)—H^+^ (with the water present in 1 case); this chain extends below the asparagine level, that is, below the gating region. The K^+^ in the gating region both enter with only 5 hydrating water molecules; the reason one water appears to be lost before entering the gate region is not entirely clear, but appears to be a consequence of interactions at the bottom of the gating region. The other K^+^ ions in the calculation, which stay below the gate, are not relevant to the question of dehydration at the gate. The upper K^+^ is near the upper boundary of the gating region, where it can enter the pore cavity; it retains only one water molecule, with two waters at approximate hydration distance strongly tied to another water, thus unable to move with the ion. The lower K^+^ that is within the gating region still retains two waters of hydration, and will replace the upper ion at that energy minimum, where it becomes the upper ion, with only one water of hydration. Each carboxyl has more than one interaction, not limited to the water that it pulls from the K^+^. Thus, in this case (P475D), the hydronium ions fail to prevent dehydration of the K^+^, leaving the ion sufficiently dehydrated to pass through the gate.

#### Summary

It is not a surprise that a system with this many components produces such a complex result, which makes it a severe test of the hypothesis that dehydration of the K^+^ is required to allow passage of the ion. Based on experimental results, the P475D mutation is constitutively open at physiologically relevant potentials, the P475S mutation is closed; the calculated results turn out to be consistent with these experiments, and to add details to the hypothesis.

In the P475D case, all protons optimized well below the aspartates, down to where the boundary of the gate region reaches the intracellular solution. In the region near the aspartates, where dehydration takes place, there is little difference with protonation, except in details that do not appear to be relevant to dehydration. The aspartates are more flexible than the prolines they replace, and at least one bends down; unlike the prolines, the aspartates do not remain in an almost planar conformation. The diagonal distance between carboxyl oxygens is 8.7 Å, 2.7 Å *less* than the corresponding distance in the non-conducting serine mutation; the distances come from the optimized structures (and although approximate, differ by much more than any possible uncertainty). This shorter distance means that the K^+^ ion can leave three of its waters with the aspartate, and a fourth water hydrogen bonded to three other water molecules; the K^+^ can move further into the pore with two water molecules, or, depending on orientation, just one. There is only one water molecule between the K^+^ and the surrounding group that can remove the water. In contrast, the serine mutation has, in all but one case, two waters separating it from the hydroxyl group, and thus five water molecules cannot be dehydrated by the serine side chain; only one water is attached to a group other than water, and could be removed. This accounts for the constitutively open behavior of the aspartate mutation, in contrast to the completely non-conducting serine mutant. Whether other anions would do as well as Cl^-^was not tested, but there is no obvious reason why this should be impossible.

The principal index of dehydration is the comparative electron density of the water molecule hydrogen to a carboxylate or water oxygen, or a Cl^-^ in some cases, vs. the water oxygen to the ion directly. The water goes with the greater electron density, which is the stronger non-covalent partial bond. The K^+^ that is entering the pore cavity has only one water attached. The K^+^ that is just entering the gating region is still mostly hydrated. In the P475D system, it is the carboxylate that helps dehydrate the ion within the region. Water – water hydrogen bonds are also critical, creating a complex network that fills the entire region. However, the upper ion (K_up_) is clearly dehydrated as it moves from the gating region to the cavity pore, which is the point that had to be tested.

## Discussion

The gate has several functions. The obvious one is to allow, or prevent, conduction of the ions through the membrane. As there is only one direction for the ions to go, this effectively reduces the degrees of freedom of ion motion from three dimensions to one. This in turn can lead to more complex consequences, but we only touch on those here. One consequence is the necessity of reducing the ion to a nearly one dimensional species, which requires the removal of most of its associated water.

We have completed a set of calculations that show that dehydration of the K^+^ ion is required for the ion to proceed through the gate region and enter the pore cavity. The calculation leads to a mechanism that prevents dehydration when extra protons are present in the wild type channel, thus blocking the passage of the ion from the gate region to the pore cavity. This does not happen in the aspartate mutation, and the serine mutation is closed even without added protons. These results are consistent with the hypothesis, but none could have been definitely asserted prior to the calculations; the aspartate and serine calculations are important because they correspond to hitherto unexplained experimental results and provide a severe test for the proton gating hypothesis. The fact that the calculations agree with the experimental results provides strong support for the validity of the first part of the hypothesis, concerning the necessity of dehydration of the ion to allow it to pass into the cavity pore.

To test the hypothesis, quantum calculations are required. While the confinement of the system by the protein is critical, the interactions that remove the water from the hydration shell of the K^+^ ion are local, involving just two atoms at a time, albeit in the context of the overall system; the polarization of the charges on the interacting groups, by the surroundings, is part of what determines the local strength of interaction. These short range interactions determine whether a water molecule remains in the K^+^ hydration shell, preventing the ion from moving forward, or transfers, either to a Cl^-^ or a protein atom, or sometimes to a strongly hydrogen bonded water molecule, allowing the ion to proceed. Determining this requires the electron density of the non-covalent interaction between the ion and the water oxygen (for K^+^) or nearest hydrogen (for Cl^-^). Classical calculations will be unable to determine these electron densities; they can only be determined quantum mechanically. Also, classical force fields generally do not allow proton transfer, which can occur here. Quantum calculations at the level used here are not perfect, but do produce a sufficiently accurate optimized structure. This structure in turn allowed a more accurate calculation to obtain its properties, in particular the non-covalent interactions among the species present.

Channel conduction is a marginal effect; the dehydration is somewhat intermittent, so that the process must be near the edge of thermodynamic stability (but see the comment about thermodynamics below); the open channel gives an oscillating (flickering) current, suggesting that the channel may be intermittently closed. Also, if closing the gate is due to protons that produce a change in the hydrogen bonding network, that network must exist close to a functional (open/close) threshold. In figures 4A and 4B we have details of the dehydration in an open channel. Dehydration requires the electric field to overcome as much energy as holds the permeant ion; the ion does not move when there is no field—that is there is no current when the field is zero, to a good approximation. This is discussed, for example, by Wang and Sun (70), and is at least consistent with the experimental rectification behavior of wild type channels, which shows an I – V curve characteristic of a rectifier. However, the Wang and Sun result is based on an MD simulation, and is not necessarily quantitatively reliable. Lynch et al (82) have provided an extensive review of water in pores, based primarily on MD simulations. However, their review does not consider the type of channel we are concerned with, and their consideration of quantum effects concludes that these were too difficult to compute with the computer resources existing up to the time of that review; for a full study of dynamics, this is still the case with respect to dynamics, although certain inferences can be derived from the static calculations which are possible now. MD with polarizable force fields shows more ability to hydrate hydrophobic surfaces than MD with non-polarizable potentials, as has been demonstrated by a study of the TMEM175 channel (83). It is clear that MD with a non-polarizable force field is unlikely to be reliable. It is also very likely that quantum effects are not negligible, as shown by the electron density in non-covalent interactions in the results presented here; also, see the discussion below concerning the de Broglie wavelength of the proton. Therefore, there is reason to doubt even classical MD with polarizable force fields.

Dehydration in wild type is marginal, as expected to account for the observed behavior, in which the open channel shows flickering conduction, and the open probability is less than 1. While adding an electric field turned out to be too difficult for us to calculate, qualitative considerations are consistent with the observed result.

The first three of the O – Cl^-^ electron density overlaps are negative in Table 2, case 2; these interactions are repulsive, so the Cl^-^ ions will not dehydrate the K^+^ ion; the presence of H^+^ completely reverses the nature of ion pairing in the gate, largely by forming solvent separated ion pairs. This cannot be deduced from the structure alone; the bonding is needed to understand the importance of the added protons. The prolines do not interact with the water at all—the electron density overlap is 0.000 in all cases for the proline nearest to the ion furthest up in the gate (closest to leaving the gate region for the pore cavity), and the optimized configuration is not symmetrical. Table 2 case 1, with the large (6.68 Å) separation of upper and lower K^+^, with the lower K^+^ at the bottom of the gating region, is the only calculation of all those we have done that is not obviously in agreement with the hypothesis. When we examine the configuration of the water, and the relation to the ion, we can see that there is a network that has formed around the ion; when it attempts to move up in response to an electric field, it will have to break several hydrogen bonds in order to advance. While further work is needed to see in detail how this is bonded (this result is not as clear cut as the rest of the calculations), the result, although introducing a complication, does not contradict the hypothesis. We have not worked out just how close the water below the upper K^+^ comes to being a bridging water to the K^+^ below; the distance (6.88Å) is very close to what is required, so it would not be surprising if part of what holds the K^+^ back is a water anchored, or bridged to, a K^+^ below. The hydration of the ion in the network may be more complex than just a question of the direct dehydration of the ion. Considering the complexity of the network it is almost a surprise that direct dehydration is sufficient in almost all cases.

### Water rotation and rearrangements

Since interactions with the water are critical, it is necessary to suggest how the water moves. The calculations we have done provide local minima without dynamics. Nevertheless, we can infer certain steps that are necessary. There will be one or two water molecules that track the ion, and therefore must rotate. In doing so, the water dipole will at one point in the rotation point in the direction that conflicts with the local electric field, especially that which is generated by the local dipoles. This is not impossible; indeed, it is analogous to the well-known Bjerrum defects found in ice. Therefore, there is no problem with the fact that some intermediate transient states must allow energetically unfavorable orientations. We do not find these in our calculations, as we obtain energy minima. There must be some energy barriers in the course of the progress of the K^+^ ion. Considering the rather limited potentials, on the order of at most a few k_B_T, that are sufficient to generate a current, the barriers must be relatively small, suggesting that the ion path must be well fitted to accommodate the passage of the ion.

Conclusion from the tables: in all open (i.e., no extra proton) cases, water is held, in many cases by chlorides (greater electron density for their non-covalent bonds), while in the closed (with four protons) cases that correspond to figure 4, K^+^ maintains its hydration, or at least, too much of it to allow progress into the pore cavity. The protons form solvent separated ion pairs with the Cl^-^, preventing those ions from removing the hydrating water; as they also affect the water network, this is not the only thing they do, but it is the most relevant to the hypothesis. (There is also competition in the selectivity filter among ions, in a different type of ion channel (84), but the relation to gating seems to be only by analogy).

The new calculations show how ions, particularly protons, at the gate of Shaker (and therefore probably K_V_1.2) can control ion movement and ion dehydration in the gate region of these channels. While we have been discussing water in regard specifically to the dehydration of ions at a channel gate, there is a very large, more general, literature, discussed in the introduction, concerning water in nanochannels, including the hydration of the ions, the formation of ion pairs and clusters, and the orientation of water in the confined spaces on the order of a nanometer or less, as discussed in the review by Zhang et al (40). A good deal of that work concerns water in carbon nanotubes, or next to a graphene surface; although the channel gate is not so hydrophobic, much of the pore beyond the gate is fairly hydrophobic, so the literature on carbon nanotubes may not be entirely irrelevant to channels. To the extent that the gate interacts with the selectivity filter at the other end of the pore (and there is some evidence that it may, at least in KcsA (85)), the cavity between may be important. However, the focus of this paper is the transition from the gate to the pore, so we do not discuss the transition onwards to the selectivity filter.

In water confined in spaces the size of ion channels, about 1 nm, the water can be aligned with the walls, the dielectric constant dropping almost to its minimum possible value around 2 (the confined water dielectric constant is not always considered to be this low, but it is not near 80, which requires free rotation), and the ion-wall interactions may dominate, while ion pairs, and even clusters, may form. The dielectric constant is a matrix and need not be the same in all directions; asymmetry allows the rotation to be maintained in a plane orthogonal to the gate axis, whereas rotation can be so restricted parallel to the axis that that component of the dielectric is much smaller. Our results suggest that the strength of the water-wall interactions is particularly important, but may be indirect, involving ions in addition to the water and the wall. Liu et al (86), consider complex interactions in a confined space; chloride, for example, behaves differently in the confined space than in bulk. Peculiar structures are possible, although not relevant here. For example, chlorides of divalent cations can form two-dimensional structures in a suitably confined space (46).

More important, based on the calculations, is the distribution of electron charge density and its interaction with the ions in the channel. This too is dependent on the orientation of water molecules in the gating region. The orientation is determined in part by the Cl^-^ and K^+^ ions, so the overall minimum energy configuration constitutes a cluster covering a substantial part of the gating region. The water, the potassium ion, the counter-ions, and the boundary, taken together, must fit. If the boundary is too wide, the cluster fails to form, and the ion does not move forward. The advance of the ion is marginal, and the electric field, although not a strong force, is sufficient to allow the ion to move, when it is dehydrated. While Brini et al (43) do not discuss channels, they show how the properties of water, as a molecule, make possible much of what we see in our calculations. In the case of the serine mutant, the ion is separated from the wall by two water molecules, for all but one water molecule that does contact the protein; at least four, probably five, out of six water molecules remain hydrating the ion, thus preventing its passage out of the gate region. The results section shows the difference between the serine and the aspartate mutations; the latter has one water between ion and carboxyl group (the relevant part of the protein in this case) instead of two, and the hydrating water can be left behind, so that the ion can advance in the aspartate case.

We have cited several relatively recent references (27–30) that considered the role of water in ion channel gating; actually, an earlier paper (77) already showed preliminary quantum calculations relevant to this topic, and still earlier work introduced the idea that water was key to gating, the earliest such proposal going back almost 40 years (87). However, the consensus in the field for at least 30 years has been that water is not very important for gating, with the opening and closing of the gate being a mechanical process in which the S4 transmembrane helix in each of four domains is pulled down by the electric field, with the S4 then somehow pushing the S4-S5 linker against the pore residues, mechanically closing the pore entrance (8). Removing the field reverses this sequence, putatively opening the channel gate. We have reviewed the evidence for this type of model (88), and found that essentially all of it is subject to alternate explanations. However, we had not previously produced evidence from our own calculations that shows how the dehydration model, with a role for protons, could work. This evidence, presented in this new set of calculations, appears to be only consistent with a gating mechanism in which water, dehydration, hydrogen bonding in confined spaces, ion pairing (including solvent-separated ion pairing), and the effect of protons, are all components of the gating mechanism. This also accounts for the fact that K^+^ arrives at the gate with six (possibly in some cases five or seven) hydrating water molecules and leaves with only two, or even one.

The generality of this mechanism is also worth considering; these calculations having been done on one type of potassium ion channel, with a particular structure, it is not necessarily the case that another type of potassium channel, let alone a sodium channel, would gate in the same manner. The protonatable residues are different. The dehydration of the sodium ion is still necessary, as the ion arrives at the gate with more water than it has when it leaves the selectivity filter, but is more difficult than that of a potassium ion. This said, there is evidence that, at least in certain bacterial sodium channels, the S4 segment does not move in channel gating, strongly suggesting that that sodium channel, and very likely others, also depend on a mechanism similar to that we propose for the potassium channel (74). It is interesting that the sodium channel gate has a section lined with glutamate. In addition, D_2_O makes a difference in gating at least sodium channels, and there are osmotic effects (53, 55), so water must be involved. Separate calculations would be needed to show how dehydration works in this case, and the details of how water is involved in gating; complications remain that are difficult even now to sort out, concerning the behavior of ions at surfaces, especially biological surfaces; in particular, the activity of water, as well as that of ions, is difficult to determine. One thing that is clear is that water cannot be ignored. The review by Menon et al does discuss this (42).

### Electron densities, and the relation to reaction energy

The electron densities require some interpretation, as we rely on these for understanding the results of the calculations. In many cases we depend on differences in these densities to decide whether a water molecule remains with the K^+^ ion, or is removed. These electron densities represent a level of weak bonding that amounts to a fraction of a covalent bond. Caution is required in interpreting the strength of the interactions (see the discussion at the end of the Methods section), but it is fairly reasonable to take the bond density to represent the fraction of a two electron covalent bond. This makes a bond density of 0.1 around 0.05 of a covalent bond strength. Typically, such a bond might be about 400 kJ, as for a C – H covalent bond, and is defined in the gas phase; however, here we are dealing with a local bond pair, so it is not unreasonable to take the covalent bond strength to be about the same as for an isolated bond. 400 kJ is about 160 k_B_T. We are dealing with electron density differences of at least 0.05. If we assume that the electron density does represent a fraction of a covalent bond strength, even 0.05 would be about 8 k_B_T, far from negligible in this context. Most of the electron density differences are about 0.1 or greater, thus around 16 k_B_T or more, possibly a rough estimate, but certainly not negligible. Even if the electron density differences are not exactly proportional to the bond strength, it is reasonable to use them as indicators of the interaction strength of a water with its neighboring groups, and with which it shares electron density. Negative electron density suggests a repulsive, or at most negligible, bonding interaction. The electron densities that we find are consistent with this interpretation. The dehydration of the K^+^ ion is endothermic, and in fact ΔG > 0 for removing the water. This tells us that the free energy gained by joining the water to the competing ion or other species must be larger or at least comparable. The energy that we expect from the electron density values, as considered above, is great enough to effect the dehydration. One hydrogen bond is generally larger than k_B_T (k_B_, Boltzmann’s constant; T, temperature, here roughly 300 K), so the energetics is reasonable.

*Osmotic effects* are particularly likely to be significant, as the ion concentration varies on a nanosecond time scale; it is not clear what time scale is relevant for the change in the activity of water, but it seems likely that it is faster than the nanoseconds for ion concentration to change; transmitting the information about the activity of water is likely to proceed through hydrogen bonds, which can rearrange on a tens of picoseconds time scale. Based on present data, including that from the calculations presented here, we can conclude that water is a central player in gating, that the potassium ion must be dehydrated in order to pass through the gating region, and that protons can prevent dehydration. How this couples to the osmotic effects that must occur as the ion concentration changes in the gate region, and to the external electric field, will require further computations. Ion concentration changes not only with the entry and exit of the current carrying ions, and the protons, but with fluctuations in the composition of the gating region, as we consider below.

### Thermodynamic and energetic considerations, other than osmosis

We suggested just above, in the electron density discussion, that we should expect small but non-negligible fractions of bond energies. However, there are additional energy questions. In effect, the optimized structures are found at 0 K, which appears to present a problem. However, the entropy contribution to hydration free energy at room temperature is only about 5% of the total free energy of hydration of the ions, so this is probably not a serious impediment to the use of the calculation (77, 78); these references give the most useful values of the hydration energy of the ions. From the low absolute value of the entropy (TΔS as a fraction of the total free energy), it is reasonable to infer that the energy term dominates the hydration free energy, and the energy minimization is therefore adequate to determine the essential behavior of water and ions in the channel gate. The differences in heat capacity of the open and closed states of the channel are also unlikely to be large, so when integrated to 300K, they are unlikely to alter the relative free energies of dehydration in the channel gate. All structural models, as well as the initial positions of the protein in these optimization calculations, are also determined, in both cryoEM and X-ray crystallography, near or below the minimum temperature of high density amorphous water, so these structures are generally the same as would be found at 0 K.

There is also a question as to the applicability of thermodynamics to gating; it is almost certain that the gating region is not at equilibrium, even when closed, so strictly speaking thermodynamics does not apply. In the open channel, there is a current with applied electric field, which by definition means that there is no equilibrium. However, it is possible to consider a gradient of ΔG between different “states” of the system, where “states”, and ΔG, are only approximately defined; while out of equilibrium, they may be close enough to use this language. This gradient acts in the direction of moving the K^+^ forward, and thus helps in understanding some of the driving force for the opening of the channel. The fluctuations in the system are so large that they constitute another reason for questioning the applicability of standard thermodynamics. Thermodynamics of Irreversible Processes may apply, as with the streaming current measurements that we have referred to. However, using thermodynamics requires great caution in a system that lacks a well defined equilibrium.

### Fluctuations

There are fluctuations comparable to the size of the gating region. The gating region volume is small enough that at bulk density, it would contain about 25 – 30 water molecules, if we ignore the ions, which also take space. If there are 25 water molecules and we expect fluctuations of the order of the square root of the number of molecules in the gating region (essentially by exchanging with the intracellular fluid), the fluctuations would amount to around 20% of the average number of water molecules present. One must expect flickering in properties. Even one ion entering the gating region, and displacing one water, makes a four percent change in water number, in addition to a possibly drastic rearrangement of the hydrogen bonding network. This means a change in the thermodynamic activity (partial molar Gibbs energy) of the water. It is hard to know how large the change is, and almost certainly both ion transit and fluctuations produce difficulty in defining the activity of the water. Therefore, we do not consider the thermodynamic properties of the overall system to be sufficiently well defined to provide a really useful description of the gating region.

### Flicker noise

If the fluctuations are important, and the open-closed transition is near a threshold, one must expect flickering in conduction, as the gate switches configurations. Flickering conduction is observed with every instance of the open channel. Some of the apparent closings are long enough that the channel may even go to a temporarily (millisecond time scale) blocked state, or some other non-conducting state that is not inactivated (conductance recovers). This suggests that an ion, or ion pair, is in a configuration in which the ion fails to lose its hydration shell, and that even though such a configuration is not stable, it lasts for long enough to appear in the current record. The conductivity itself suggests that the energy changes are small between conducting and non-conducting conditions. Continuing the discussion about energy from the electron density section: A change in potential of the order of k_B_T/q ≈ 25 mV; (q=1.60 x 10 ^-19^ coulombs) effects a significant increase in current. Here we do not try to quantitatively compare this conductivity to the energy differences that apply to the complete system; local energies, or the distribution of the field, would be required for this, but are not available. The fact that we observe electron densities in non-covalent bonds on the order of 0.1 suggests that we should expect voltages that correspond to energy differences on the order of k_B_T, as is observed. We should not be surprised that the difference between a fully conducting state and a state in which an ion cluster momentarily blocks the channel is small and fluctuations in position and orientation of the water, and position of the ions, can make the difference between momentary conducting or non-conducting configurations. Water rearrangements under osmotic gradients would couple to these effects. Taken together, these effects could account for flicker noise. Finding direct evidence for this proposal remains for future work (the nanocavity at the exit of the selectivity filter, with its digital occupancy, studied by Sumikama and Oiki (13), might also contribute to flicker noise by an entirely different mechanism). The large effect of small voltages of order of magnitude k_B_T/q suggests that flickering must be primarily derived from conditions at the gate. These considerations show that with a hydrogen bond energy being generally larger than 5 k_B_T, but with variations larger than k_B_T, ignoring switches in hydrogen bonding must lead to an incorrect description of what is happening at the gate. A strictly protein structural description of gating, without including the effect of water, especially as the ion passes through and rearranges the hydrogen bonding as it goes, cannot account for several experimental findings, among which is the fact that potentials of the order of k_B_T/q control the average current, but this current fluctuates in a manner that is hard to understand if the structure of the protein provides a stable open state..

### Topological properties, and resonance

We have not put the gating details problem completely to rest with these calculations. One possibly relevant topic we have not considered are the topological properties of the hydrogen bond arrangement in the confined space of the gate. The topology almost certainly affects some of the energy terms. The question for water in a confined space, as well as the arrangements of water in hydrogen bonding cycles and related topological properties, has been addressed by several groups (89–93). Including the effects of protons is necessary. Rings of bonds that allow hydrogen bond transfer may produce resonance effects; there is an example in which a ring containing phosphate and arginine shows a definite resonance effect (94). Resonance effects are necessarily included in quantum calculations, although some revisions may be produced by more precise calculations. However, there is not much known about topological properties of hydrogen bond networks with ions present, at least in confined spaces. At the gate, the presence of protons will force rearrangements of cycles (or graphs) of hydrogen bonds, thus producing different properties. Further work will be required to provide a complete picture of the way the ion proceeds through the gate, although the calculations presented here clearly provide the framework of what must happen. As the ion moves, the hydrogen bonds in the gate will rearrange. There is no question that the rearrangements of water are critical for this process; to compare channels, understand the effects of mutations, and possible drugs for the channels, will require working out the topological details of the steps as the ion progresses through the gate.

As hydrogen bond rearrangements are inevitable when an ion passes through the confined water at the gate, and these involve energies that are comparable to the energy derived from the applied field that generates the measured current, or larger, ignoring these rearrangements is almost certainly an oversimplification. Understanding the interaction between ions and water, and the effects of water rearrangements, is obviously critical to understanding channel gating. We have, with these calculations, established the framework for understanding channel gating. Additional work remains on details of the hydrogen bonding rearrangements, together with the dynamics of the ion transport, and specific non-covalent interactions, as well as quantitative understanding of energetics.

## Conclusions

1. Water is stripped off the hydration shell of a potassium ion at the gate of an open channel, at least for the Shaker channel, the channel computed here. The open channel can pass only one or two of the original 6 (sometimes 5) waters of hydration. Competition for water from the ion hydration shell among ions (especially counterions), protein side chains, and strongly hydrogen bonded water molecules, is a major mechanism for accomplishing this. The results are sufficient to support the proton gating hypothesis, even if by a relatively small margin in some cases. The fact that the margin is small is consistent with observations: large fluctuations in current, and current dependence on weak electric fields. This consistency increases confidence in this overall framework for gating.
2. Mutations in the protein wall of the gate may affect local interactions. These local, or short range, interactions are seen here as the non-covalent electron density linking neighboring atoms or ions, a form of partial bonding. Although these are local interactions, they are calculated in the presence of the complete system, so that they have the effects of the entire system included in determining the strength of the local interactions, which show how strongly the water molecules involved are interacting with the K^+^ ion and its neighbors. They include the effect of polarization of the molecules, and electron transfer amounting to partial bonding, in the context of the entire system.
3. Our hypothesis is that the presence of excess protons in the closed state of the wild type channel prevents dehydration of the ion, thereby preventing the ion from advancing out of the gate into the pore cavity. Previous work, cited above, showed how these protons would come from the voltage sensing domains, and constitute gating current. Here we see how they alter the water network in the gate, preventing enough dehydration to allow passage of the K^+^ ion, thus closing the gate.
4. In the serine mutant, there are at least four waters that remain hydrating the ion. There is nothing that might compete for that water; all other hydrating waters on the K^+^ are separated by one more water from competing groups, so that these waters are not removed; the ion cannot proceed further, making the channel effectively closed. Possible connection to second shell water might impede ion progress further. This is confirmed by electron densities. With the aspartate mutation, this is not the case; the hydrating waters can be removed even when protons are present, and the ion is therefore able to progress, so that the channel is effectively constitutively open.
5. Further work is required to determine: *i*) applicability of this mechanism to other channels, as well as some details of the water-ion interactions determined by a higher level quantum calculation *ii)* checking exactly how the dehydration proceeds when the two K^+^ that reach the gate region interact, and what the significance of the distance between them in the optimized structures might be *iii)* details of interactions with the external electric field, as well as osmotic effects *iv*) additional quantum corrections that might be found by not using the Born-Oppenheimer approximation for protons *v*) resonance effects from rings formed by hydrogen bonded water molecules are necessarily included in these calculations, but dynamics of how the topology of the water plus ion networks that form in the gating region is disrupted by added protons remains to be elucidated. *vi)* Fluctuations also affect osmotic activity, resonance effects, and hydrogen bond topology, and remain to be calculated. However, the fact that gating occurs near a threshold of transition between open and closed conditions can be understood from these calculations.

## AUTHOR CONTRIBUTIONS

AMK and MEG: Conceptualization of problem, planning research, carrying out research, evaluation of results, and writing. RRM: Writing, editing, evaluation of results.

## ACKNOWLEDGEMENT

The authors acknowledge the Texas Advanced Computing Center (TACC) at The University of Texas at Austin for providing the computational resources on the Stampede3 computer that produced the results reported in this paper. URL: http://www.tacc.utexas.edu

